# Cellular Heterogeneity of Pluripotent Stem Cell Derived Cardiomyocyte Grafts is Mechanistically Linked to Treatable Arrhythmias

**DOI:** 10.1101/2022.09.15.500719

**Authors:** Dinesh Selvakumar, Zoe E. Clayton, Andrew Prowse, Steve Dingwall, Jacob George, Haisam Shah, Siqi Chen, Robert D. Hume, Laurentius Tjahjadi, Sindhu Igoor, Rhys J.P. Skelton, Alfred Hing, Hugh Paterson, Sheryl L. Foster, Lachlan Pearson, Emma Wilkie, Prajith Jeyaprakash, Zhixuan Wu, Jeffrey R. McArthur, Tony Barry, Juntang Lu, Vu Tran, Richard Bennett, Yasuhito Kotake, Timothy Campbell, Samual Turnbull, Quan Nguyen, Guiyan Ni, Stuart M. Grieve, Nathan J. Palpant, Faraz Pathan, Eddy Kizana, Saurabh Kumar, Peter P. Gray, James J.H. Chong

**Author notes:** Corresponding author: A/Prof James Chong, The Westmead Institute for Medical Research, 176 Hawkesbury Road, Westmead, Sydney NSW, 2145, Australia. Contributed equally to this work (Dinesh Selvakumar, Zoe E. Clayton). Declarations of Interest: None.

## Abstract

**Background:** Exciting pre-clinical data have confirmed that human pluripotent stem cell derived cardiomyocytes (PSC-CMs) can remuscularise the injured or diseased heart, with several clinical trials now in planning or recruitment stages worldwide. However, ventricular arrhythmias are a predictable complication following engraftment of intramyocardially injected PSC-CMs. Therefore, there is an urgent unmet need to gain mechanistic insights and treatment strategies to control or prevent these engraftment arrhythmias (EAs).

**Methods:** We used a porcine model of myocardial infarction and PSC-CM transplantation to investigate efficacy of pharmacologic and catheter based anti-arrhythmic strategies in mitigating EAs. Furthermore, cell doses were robustly phenotyped using single cell ribonucleic acid sequencing and high parameter flow cytometry to identify cellular characteristics predictive of arrhythmogenesis.

**Results:** Combination therapy with amiodarone and ivabradine significantly reduced EA rate and burden following PSC-CM transplantation. Catheter ablation was also a feasible and effective treatment strategy which could be considered in the case of pharmacologically refractory arrhythmias. In addition, we show that EAs are mechanistically linked to cellular heterogeneity in the input PSC-CM and resultant graft. Specifically, we identify atrial and pacemaker-like cardiomyocytes as culprit arrhythmogenic subpopulations. We further describe two unique surface marker signatures, SIRPA^+^/CD90^-^/CD200^+^ and SIRPA^+^/CD90^-^/CD200^-^, which identify arrhythmogenic and non-arrhythmogenic cardiomyocytes respectively.

**Conclusion:** Our data deepens mechanistic understanding of EAs and suggests that modifications to current PSC-CM production and/or selection protocols could ameliorate this problem. We further show that current clinical pharmacologic and interventional anti-arrhythmic strategies can control and potentially abolish these arrhythmias, an important safety consideration given several impending clinical trials.

## Introduction

Myocardial infarction (MI), the leading cause of heart failure, results in the loss of up to 1 billion, highly specialised cardiomyocytes (CMs)^1^. Despite great interest, numerous attempts to replace damaged or destroyed cardiomyocytes with cell therapy has yielded inconsistent results in clinical trials^2–4^. This may partly be attributable to the lack of cardiomyogenic differentiation capacity of tested cell types^5, 6^. Pluripotent stem cells (PSC) however, are a renewable source of CMs^7^. Exciting pre-clinical data have confirmed that PSC-CMs can remuscularise and improve cardiac function in clinically relevant large animal models^8–13^. Several clinical trials are now in planning or recruitment stages worldwide^14–18^. However, hurdles to widescale clinical translation remain^19, 20^. The most concerning of these relates to ventricular arrythmias which ensue following intramyocardial PSC-CM delivery, hereafter referred to as engraftment arrhythmias (EA)^8–12^.

There is a paucity of literature examining the mechanism of EA, with existing evidence suggesting arrhythmias arise due to the abnormal impulse generation and enhanced automaticity of PSC-CM graft^9, 11^. Furthermore, there has only been one study on therapeutic strategies to mitigate EAs^12^, with this recent report suggesting that pharmacotherapy can suppress but not abolish arrhythmias. Therefore, as PSC-CMs enter the clinical arena, there is an urgent need to gain further mechanistic insights and test additional means of therapeutic arrhythmia control.

Current PSC-CM differentiation protocols yield heterogeneous cellular outputs. Although ventricular cardiomyocytes predominate, other cell types such as atrial and pacemaker-like cells are also present^21–23^. Despite suggestions that purified ventricular cardiomyocytes may be the most desirable cell product for transplantation applications^24^, to date no study has investigated the important relationship between cellular heterogeneity and arrhythmogenesis. Here, we use a porcine MI model to study EAs after phenotyping the composition of PSC-CM cell doses. We aimed to identify cellular characteristics predictive of arrhythmogenesis, hypothesising this would inform the safe production of cardiomyocytes for future clinical use. We also sought to test efficacy of clinically available pharmacologic and procedural anti-arrhythmic treatments that could abrogate EAs should they arise in clinical trials.

## Methods

Further details of all experimental methods can be found in the supplementary material.

### Cell Production

The H9 cell line containing the gCaMP6f fluorescent calcium reporter was used for all transplantation experiments. Cells were expanded and differentiated in a stirred-tank bioreactor system using a small molecule differentiation protocol (Supplementary Fig. 1a)

### Single-Cell RNA Sequencing (scRNa-seq)

PSC-CMs were hash tagged for scRNA-seq. The cell suspensions were loaded onto Single Cell 3’ Chips (10x Genomics) to form single cell gel beads in emulsion. The Chromium library was generated and sequenced on a NextSeq 500 instrument (Illumina).

### High Parameter Flow Cytometry

Thawed PSC-CMs were stained with an amine reactive viability dye and fluorochrome-conjugated membrane marker antibodies (Supplementary Table 4). PSC-CMs and compensation controls were analysed with a FACSymphony A5 cytometer (BD) and FACSDiva software (BD). Data analysis was performed using FlowJo software (BD) (Supplementary Fig. 2).

### *In vitro* Electrophysiology

Patch clamping and optical electrophysiology experiments were conducted using iPSC-CMs (SCVI-8, Stanford Cardiovascular Institute) on Syncropatch 384PE (Nanion Technologies) and Kinetic Imaging Cytometry (Vala Sciences) instruments.

### Porcine Experiments

Porcine experiments were conducted in landrace swine weighing 25-30kg. Experimental protocols (Fig 1a) were approved by the Western Sydney Local Health District Animal Ethics Committee. All animals received humane care in compliance with the Australian National Health and Medical Research Council guidelines.

**Figure 1.**
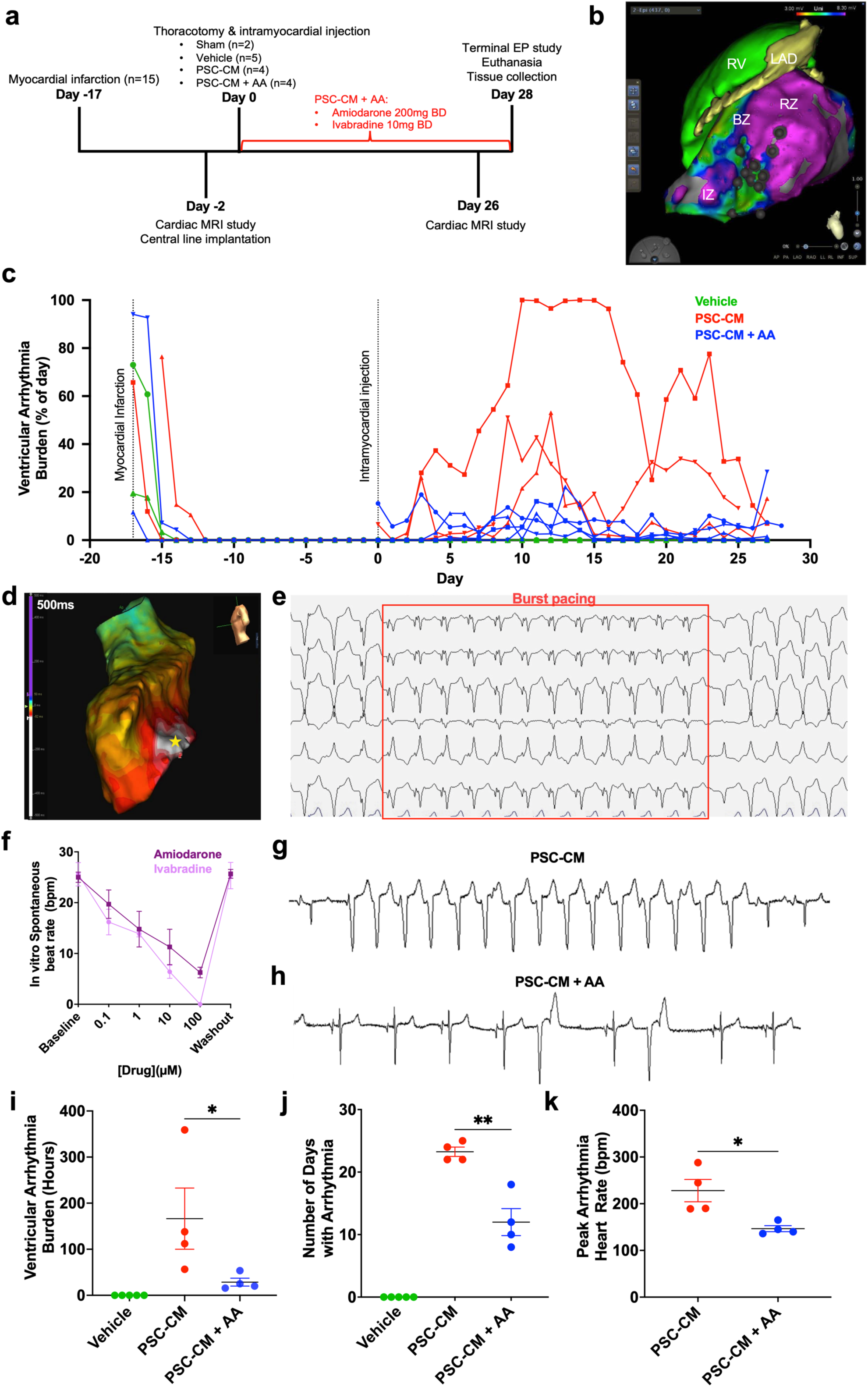
Focal, automatic engraftment arrhythmias can be effectively supressed with pharmacotherapy. **(a)** Study timeline for phase 1 large animal experiments in which 13 swine were randomised to 4 treatment groups (PSC-CM, PSC-CM + AA, Vehicle) following percutaneously induced myocardial infarction. A further 2 animals underwent sham infarction and vehicle injection. **(b)** Representative CMR reconstruction (created with ADAS 3D) merged with epicardial voltage map, used to choose, and annotate border and infarct zone injection sites (black circles). **(c)** Plot depicting the percentage of time per day each cell (red), cell and drug (blue), or vehicle (green) recipient spent in ventricular arrhythmia over course of study protocol. **(d)** Representative endocardial activation map during EA localising arrhythmia origin to focal sites of cell injections (white: early activation, purple: late activation, star: site of earliest activation) **(e)** Representative EA persisting through burst pacing suggesting an automatic rather than re-entrant mechanism. **(f)** Dose-dependent suppression of PSC-CM contraction rate using amiodarone (dark purple) and ivabradine (light purple) *in vitro* (0-100µM drug concentrations). **(g)** Telemetry strip from representative cell recipient exhibiting run of EA with spontaneous onset and offset. **(h)** Telemetry strip from representative cell and anti-arrhythmic recipient showing predominantly sinus rhythm with occasional ventricular ectopic beat. **(i-k)** Arrythmia parameters from day 0 to day 28 for each subject, grouped by treatment allocation (presented as mean ± SEM). Significant reduction in **(i)** total hours of arrythmia (*p = 0.03, Mann Whitney test), **(j)** days with arrhythmia (** p = 0.003, unpaired *t*-test, df = 6), and (**k)** peak arrhythmia heart rate (* p = 0.02, unpaired *t*-test, df = 6) in cell recipients treated with anti-arrhythmics.

#### Myocardial Infarction

MI was percutaneously induced 17 days prior to cell injection. The left anterior descending artery was occluded distal to the first diagonal branch with an inflated angioplasty balloon for 90 minutes.

#### Telemetry device implantation and analysis

ECG and accelerometer data was continuously monitored from a subcutaneously implanted telemetry unit (EasyTEL+, Emka Technologies). Semi-automated quantification of recorded data was performed offline using the device specific software package (ECGAuto, Emka Technologies)

#### Cardiac Magnetic Resonance Imaging (CMR)

CMR was performed on a 3 Tesla Prisma system (Siemens).

#### ADAS-3D CMR Reconstructions

ADAS-3D (Galgos Medical), a CMR segmentation software, was used to generate three dimensional reconstructions of the left ventricle for integration with the electroanatomic mapping (EAM) system.

#### Thoracotomy, epicardial mapping and cell injection

The epicardial surface of the left ventricle was electroanatomically mapped after exposure through lateral thoracotomy. The generated voltage map was used to identify scar, border, and remote zones. ADAS-3D reconstructions were imported to the CARTO EAM system (Biosense Webster). Injections of either vehicle (8x 300 µL injections of cellular media) or PSC-CMs (750x10^6^ cells distributed in 8x 300 µL injections) into infarct and border zones were performed using a 27-gauge insulin syringe.

#### Immunosuppression

Immunosuppression was administered to prevent xenograft rejection. Starting 5 days prior to transepicardial injections until euthanasia, oral cyclosporine A (10-15mg/kg) was given twice daily maintaining trough levels greater than 250ng/mL On the day of injections, 500mg Abatacept, 30mg/kg methylprednisone and 5mg/kg cyclosporine A was administered intravenously (IV). From day 1 post injections until euthanasia, 100mg methylprednisone was given IV daily.

#### Anti-arrhythmic treatment

The animals randomised to anti-arrhythmic treatment received a 150mg IV amiodarone bolus at the time of cell injection. From day 1 post injections until euthanasia, they received 200mg amiodarone and 10mg ivabradine orally twice daily.

#### Electrophysiological Study

Prior to euthanasia, all subjects underwent electrophysiological studies to determine inducibility, mechanism, and electroanatomic origin of any ventricular arrhythmia using the Ensite Precision mapping system (Abbott Medical).

#### Catheter Ablation

All subjects undergoing catheter ablation were in spontaneous engraftment arrhythmia at the time of the ablation procedure. At sites of earliest activation, radiofrequency ablation lesions were delivered using a 4mm tip open-irrigation catheter (Flexability, Abbott Medical). Ablations were performed in power control mode using 30-40W of power and irrigation with normal saline. Delivery of each lesion was attempted for 30-60 seconds unless terminated prematurely due to catheter movement or impedance rise. Ablations were terminated after sinus rhythm was restored, followed by attempts to induce ventricular tachycardia.

#### Euthanasia and tissue harvest

Animals were euthanised with potassium chloride (75-150mg/kg) under deep anaesthesia. Hearts were excised and fixed in 10% neutral buffered formalin for subsequent analysis.

### Histology

Immunohistochemistry was performed on sectioned, formalin fixed hearts using an antibody panel designed to interrogate graft composition (Supplementary Table 8).

### Statistical analyses

Continuous data are presented as mean ± standard error of mean (SEM). Normality was assessed using the Shapiro-Wilk test. Statistical comparisons of normally distributed data were conducted using unpaired t tests or ordinary one-way ANOVA followed by the Sidak’s post hoc test to adjust for multiple comparisons. Non-normal data was statistically compared using the Mann-Whitney or Kruskal-Wallis test followed by Dunn’s test to adjust for multiple comparisons. Correlations were expressed using Pearson’s or Spearman’s correlation coefficients. Inter- and intra-observer CMR analysis variability was expressed using Bland-Altman plots. Survival analyses were performed using the Kaplan-Meier method, and the log-rank test was applied to determine significance between overall survival between groups. *P* values <0.05 were considered statistically significant. All analyses were performed using GraphPad Prism Version 9.3.1 software.

## Results

### Bioreactor production of PSC-CMs

Each batch of cardiomyocytes were generated from the same working cell bank material. Assessment of cardiac troponin T (cTnT) expression demonstrated an average of >80% cTnT^+^ cells and evidence of cytoskeletal formation (Supp Fig. 1d-e). Genes for pluripotency, early mesoderm induction, and cardiac specification, were assessed via quantitative polymerase chain reaction (qPCR) over the time course of differentiation (Supp Fig. 1f-i). These exhibited expression trends comparable to previously published PSC-CM production methods^25^.

### PSC-CM related engraftment arrhythmias are focal and automatic in nature

After generation of sufficient PSC-CMs, we conducted transplantation experiments over 3 phases in 23 landrace swine. Of these, 2 subjects died following induction of MI and 1 died due to procedural complications following thoracotomy.

The first phase of our large animal experiments was designed to gain insights into the electrophysiological nature of PSC-CM related EAs and to assess the treatment efficacy of clinically available anti-arrhythmic agents (AA). Transplantation studies were conducted in 15 subjects 2 weeks following percutaneously induced ischaemia-reperfusion MI (Fig. 1a). Animals were randomised into 1 of 4 treatment groups: PSC-CM (n=4), PSC-CM + AA (n=4), vehicle (n=5), or sham MI with vehicle injection (n=2). 750 million PSC-CMs or vehicle were delivered into infarct and border zones via transepicardial injections following lateral thoracotomy. To enable accurate targeting and annotation of cell injections, we developed a novel technique in which epicardial voltage maps were merged with reconstructed CMR generated from the ADAS 3D imaging software (Fig. 1b, Supplementary Fig. 3). No spontaneous arrhythmias were observed in any vehicle treated subjects, however all cell treated subjects developed EA within a week of PSC-CM transplantation (Fig 1c). Follow-up electroanatomic mapping studies performed 4 weeks post-transplantation localised EA origin to focal sites of cell injection (Fig. 1d). Subsequent histological analyses confirmed these to be sites of cell engraftment.

In the same terminal mapping procedure, EA mechanism was elucidated. All EAs occurred either spontaneously or after administration of the catecholaminergic drug isoprenaline. None occurred after programmed electrical stimulation. Termination was spontaneous, typically followed by spontaneous re-initiation. Sinus rhythm could not be restored by either rapid ventricular pacing (Fig. 1e) nor external cardioversion. Cycle length variation was noted without changes in ECG morphology. Attempts at entrainment accelerated the arrhythmia, without instances of constant or progressive fusion nor tachycardia reset. Together, these findings indicate enhanced automaticity as EA mechanism.

In contrast, vehicle treated animals only experienced arrhythmias which could be induced by rapid ventricular pacing, had fixed cycle lengths and were able to be terminated by rapid pacing or external cardioversion. These arrhythmias could be entrained from multiple right ventricular sites, leading to constant or progressive fusion along with tachycardia reset and continuation following the final entrained pacing beat. This suggests scar-mediated re-entrant circuits as the mechanism for induced arrhythmias in vehicle treated animals^26^, the typical cause of post-MI ventricular arrhythmia^27^.

### Effective pharmacological suppression of PSC-CM related engraftment arrhythmias

Given the enhanced automaticity mechanism for PSC-CM EAs, we hypothesised that suppressing the rate of graft automaticity below that of the native sinus node would reduce arrhythmia burden.

Ivabradine, a selective inhibitor of the pacemaker current (I_f_) responsible for cardiomyocyte automaticity, and amiodarone, a widely used anti-arrhythmic drug which blocks multiple ion channels, were selected as the anti-arrhythmic drugs. We hypothesised the distinct mechanisms of action for each of these drugs would offer effective rate and rhythm control in treatment of the focal automatic EAs. *In-vitro* experiments confirmed that both these drugs induced dose-dependent reduction in the spontaneous beat rate of PSC-CMs cultured in monolayer (Fig 1f). *In-vivo* arrhythmia detection was performed through blinded analysis of continuous ECG data transmitted from implanted telemetry units. Although all animals experienced transient arrhythmias attributable to reperfusion injury immediately following MI, sinus rhythm was maintained for several days leading up to epicardial injections (Fig. 1c). Amiodarone-ivabradine treatment was effective in supressing EAs with clinically and statistically significant reduction of all telemetry parameters observed in drug treated pigs (Fig. 1g-k) (Supplementary Fig. 4): total hours of arrhythmia (166.4 ± 66.4 vs 28.6 ± 8.6; p < 0.05), days with arrhythmia (23.3 ± 0.8 vs 12.0 ± 2.2; p < 0.005) and peak arrhythmia heart rate (228.0 ± 23.9 beats per minute (b.p.m) vs 146.5 ± 6.4 b.p.m; p < 0.05).

### PSC-CM with anti-arrhythmic pharmacological therapy improves left ventricular function post-myocardial infarction

Although the primary goal of this study was not to demonstrate salutary effects with PSC-CM therapy, we nonetheless assessed cardiac structure and function with serial CMR in cell and vehicle recipients. All subjects underwent CMR 2 days prior to and 4 weeks after transepicardial PSC-CM injections (Fig. 2a). Image analysis was conducted according to standard reporting guidelines by 2 blinded observers^28^ with excellent inter- and intra-observer variability (Supplementary Fig. 5). Three animals were excluded from functional analysis due to failure of infarct creation, all with scar sizes under 1.5% of left ventricular mass (Supplementary Table 2). Left ventricular ejection fraction (LVEF) was preserved in sham subjects (58 ± 1%) and similarly depressed in all infarcted animals prior to transepicardial injection (Vehicle: 43 ± 2%, PSC-CM: 38 ± 3%, PSC-CM + AA: 38 ± 6%; p = 0.44) (Supplementary Table 2). At 4-weeks follow-up, no significant change in scar size, expressed as a percentage of total left ventricular mass was demonstrated between groups (change in scar size - Vehicle: 2 ± 1%, PSC-CM: -0.6 ± 2%, PSC-CM + AA: 0.8 ± 2%; p < 0.05; p = 0.99) (Fig. 2b). Despite this, a statistically significant improvement in LVEF was observed (Change in LVEF - Vehicle: 0.3 ± 0.3%, PSC-CM: 4 ± 2%, PSC-CM + AA: 8 ± 2%; p < 0.05), with post-hoc analysis showing that this was driven by the PSC-CM + AA group (Fig. 2c-d). To further interrogate this finding, we evaluated the impact of our interventions on left ventricular volumes. Despite improvement in stroke volume (LVSV) in the PSC-CM + AA subjects (Change in LVSV - Vehicle: 10 ± 2mL, PSC-CM: 13 ± 1mL, PSC-CM + AA: 24 ± 4mL; p < 0.05), change in left ventricular end-diastolic volumes (LVEDV) were comparable between groups, and if anything, greater in PSC-CM + AA animals (Change in LVEDV - Vehicle: 21 ± 4mL, PSC-CM: 21 ± 3mL, PSC-CM + AA: 35 ± 13mL; p = 0.38, ns) (Fig. 2e-f). Together, these data suggest the improvement in function observed in PSC-CM recipients may be attributable to greater left ventricular contractility rather than reduction of adverse remodelling and left ventricular dilation post-MI, with the greatest benefit evident in animals with arrhythmias supressed by drug therapy.

**Figure 2.**
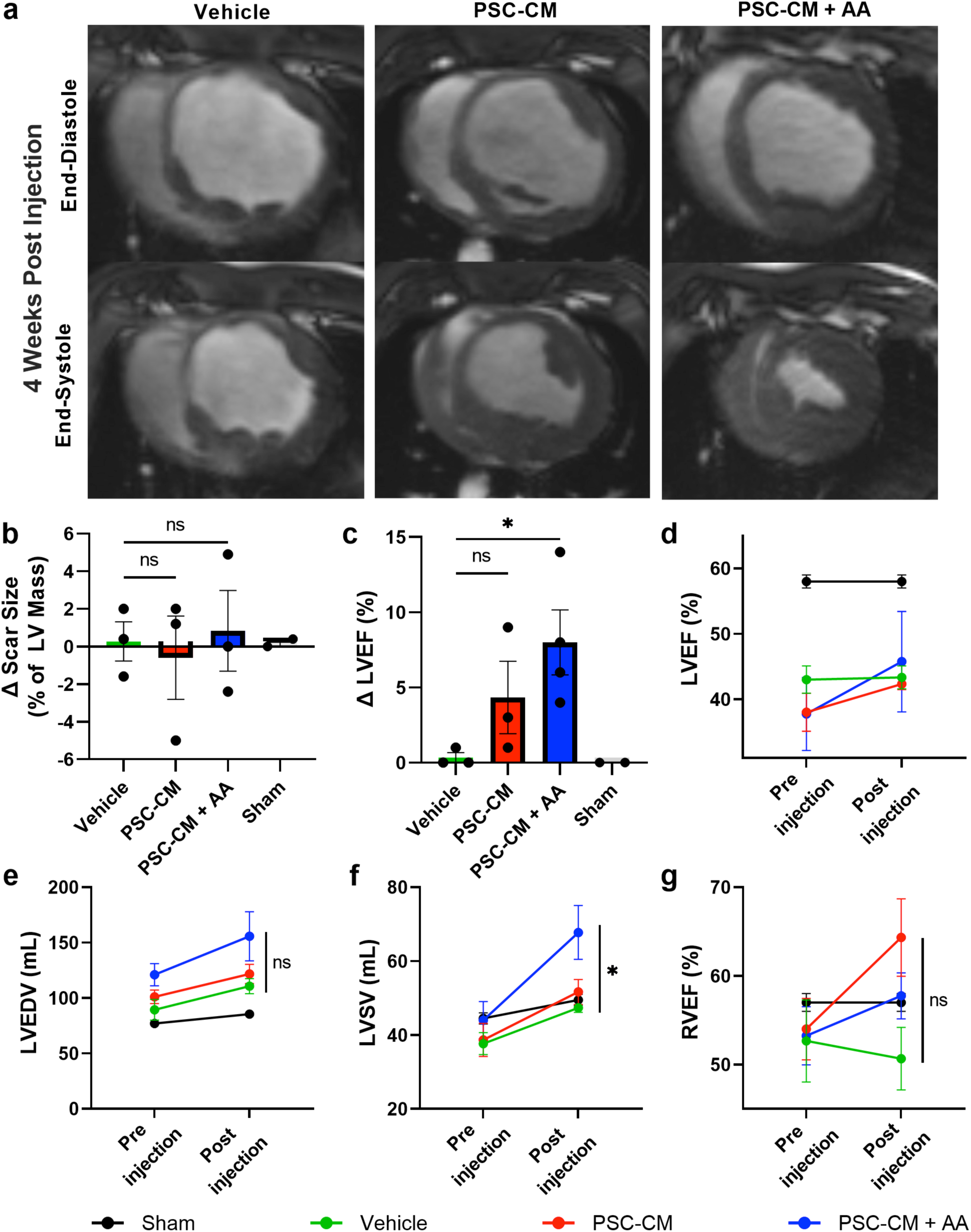
Effects of PSC-CM transplantation on cardiac volumes and function. **(a)** Representative short-axis cine MRI from end-diastolic and end-systolic phases of the cardiac cycle 4 weeks following cell or vehicle injection. Greater ejection of blood in systole for cell compared to vehicle recipients most notably in cell recipients treated with anti-arrhythmic drugs. **(b)** Plot depicting the change in scar size from baseline (post-MI, pre-injection) to 4-week post-injection, with no significant difference between treatment groups (p > 0.99, Kruskal-Wallis test) **(c)** Plot depicting the change in LVEF from baseline (post-MI, pre-injection) to 4-week post-injection, with significantly improved LVEF in cell and anti-arrhythmic recipients (Vehicle vs PSC-CM: p = 0.27; Vehicle vs PSC-CM + AA: * p = 0.03; Kruskal-Wallis with Dunn’s multiple comparisons test). **(d-g)** Pooled group data of left and right ventricular function at baseline (post-MI, pre-injection) and 4 weeks post-injection. Greatest improvement in **(d)** LVEF was noted in cell and anti-arrhythmic recipients. No significant change in **(e)** LVEDV observed between groups (p = 0.38, Kruskal-Wallis test) though there was significant improvement in **(f)** LVSV in the cell and anti-arrhythmic recipients (Vehicle vs PSC-CM: p= 0.69; Vehicle vs PSC-CM + AA: * p = 0.03; Kruskal-Wallis with Dunn’s multiple comparisons test). For **(g)** RVEF there was no significant difference between groups however a trend to improvement with cell recipients (p = 0.07, Kruskal-Wallis Test)

Therapeutic efficacy on right ventricular function was also assessed (Fig. 2g). Though a trend to improvement was evident in all cell recipients, this difference approached but did not reach statistical significance (Change in RVEF - Vehicle: -2 ± 4%, PSC-CM: 10 ± 3%, PSC-CM ± AA: 5 ± 3%; p = 0.08, ns).

### PSC-CM cell doses are heterogeneous with arrhythmogenic subpopulations

We next sought to gain phenotypic insights into input PSC-CM composition to identify arrhythmogenic cellular characteristics that could be strategically targeted. Representative samples from each PSC-CM recipient’s cell dose were retained prior to transplantation for use in scRNA-seq and high dimensional flow cytometry experiments. Uniform manifold approximation and projection (UMAP) plots of clustered scRNA-seq data identified 10 distinct cellular subpopulations, the identities of which were surmised based on differential gene expression (Fig. 3a-c). The majority of cells expressed markers of a committed cardiac lineage such as *NKX2-5, SIRPA* and *TNNI1*, representing clusters 0 to 4. Within these cardiomyocyte populations, further heterogeneity was noted. Compact ventricular markers^29–31^ such as *MYL2*, *IRX4*, *MYH7* and *HEY2* were abundantly expressed in cluster 0 whereas atrial and pacemaker markers^32–34^ including *SHOX2*, *VSNL1*, *NPPA* and *NR2F1* were localised to cluster 1.

**Figure 3.**
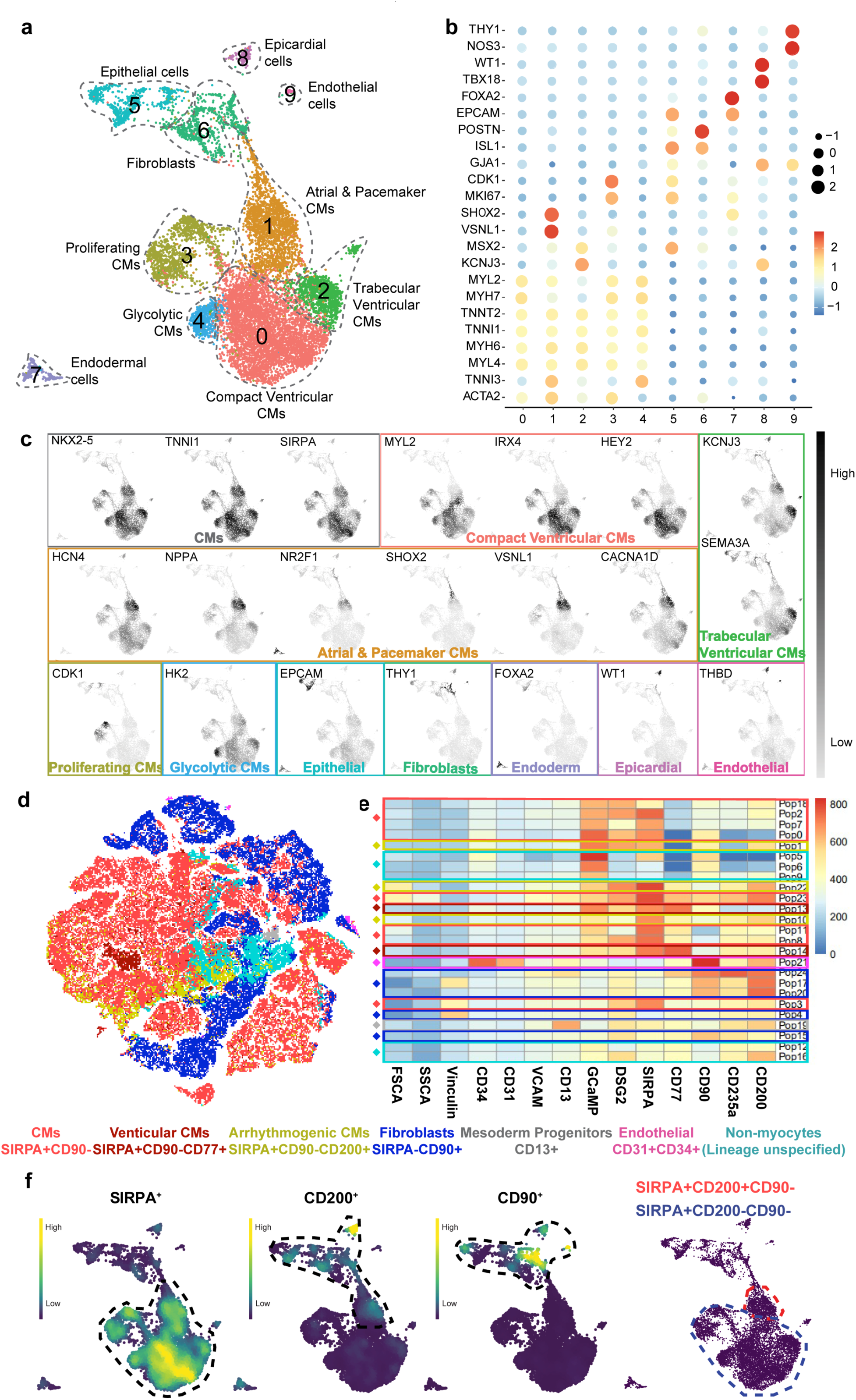
PSC-CM cell doses are heterogeneous with arrhythmogenic subpopulations. **(a)** UMAP embedding of scRNA-seq data from representative samples of cell doses. 10 clusters were identified and annotated based on differential gene expression. **(b)** Dot plot showing expression of representative marker genes used to categorise clusters. **(c)** Nebulosa plots showing the density of specific marker gene expression based on gene-weighted density estimation, demarcating cell type identity to clusters.**(d)** Representative tSNE plot from sample of a cell dose analysed by flow cytometry. FlowSOM metaclusters overlaid onto the tSNE highlight several discrete subpopulations within the cell fraction which were identified based on surface marker signatures. **(e)** Heat map showing relative expression of cell surface markers within each metacluster. **(f)** Nebulosa plots displaying expression levels of SIRPA, CD200 and CD90. Right-most panel outlining SIRPA^+^CD90^-^ CD200^-^ cardiomyocytes which were negatively associated with arrhythmias, and SIRPA^+^CD90^-^ CD200^+^ cardiomyocytes which were positively associated with arrhythmias. Arrhythmogenic SIRPA^+^CD90^-^CD200^+^ cardiomyocytes isolate to atrial and pacemaker cardiomyocytes (cluster 1) and non-arrhythmogenic SIRPA^+^CD90^-^CD200^-^ cardiomyocytes encompass all remaining cardiomyocyte clusters (clusters 0, 2, 3, 4).

Markers of trabecular myocardium such as *KCNJ3*, *SEMA3A*, *IRX3* and *SCN5A*^29, 35^ were predominantly expressed in cluster 2, with proliferative markers^36^ such as *CDK1* and *MKI67* identifying cluster 3. Cluster 4 showed strong expression of the glycolytic marker *HK2*^37^, demarcating an earlier stage glycolytic cardiomyocyte population. Non-cardiomyocyte cell populations were also present with clusters 5-9 inclusive of fibroblasts, epithelial, endodermal, epicardial and endothelial cells. Together, this data confirms the dynamic transcriptional heterogeneity of profiled cells that were used in the described animal studies.

Cell dose characterisation by way of high-parameter flow cytometry also confirmed cellular heterogeneity (Fig 3d-e). We designed an antibody panel comprising of 12 surface markers to interrogate cell composition of the PSC-CMs delivered (Supplementary Table 1). Resultant t-distributed stochastic neighbour embedding (tSNE) plots overlaid with 25 FlowSOM meta-clusters were annotated where possible based on previously reported surface marker signatures. Of note, cardiomyocytes were classified as SIRPA^+^/CD90*^-^*, with SIRPA^+^/CD90^-^/CD77^+^ cells considered committed ventricular cardiomyocytes^38, 39^. Non-myocyte subpopulations were also present, with fibroblasts defined as SIRPA^-^/CD90^+40, 41^ and endothelial cells defined as CD34^+^/CD31*^+^* ^42^. A small sub-population of CD13^+^ non-myocytes was identified, which may represent mesodermal progenitors^43^. Due to the limitations of the surface marker panel in determining cellular fate several subpopulations could not be definitively labelled and were annotated as lineage unspecified cells. Results of single-cell RNA sequencing (Fig 3a) indicate these populations likely represent cells from epithelial and endodermal lineages.

To compare the relative abundance of specific subpopulations within and between cell doses, data from all doses were concatenated and analysed. This allowed subpopulation quantification which was then correlated with total arrhythmia burden for each cell recipient (Supplementary Fig. 6, Supplementary Table 3). PSC-CM + AA subjects were excluded from this analysis due to the confounding effect of arrhythmia suppression. Interestingly, the strongest correlation between subpopulation quantification and arrhythmia burden occurred with a previously undefined SIRPA^+^/CD90^-^/CD200^+^ population (r=0.80). Conversely, SIRPA^+^/CD90^-^/CD200^-^ cells held a negative association with arrhythmia burden (r=-0.77). To further define this surface marker signature scRNA-seq data was interrogated. SIRPA^+^/CD90^-^/CD200^+^ cells isolated to cluster 1 (atrial and pacemaker cardiomyocytes) and SIRPA^+^/CD90^-^/CD200^-^ to all remaining cardiomyocyte clusters (Fig. 3f). Taken together, these data confirm the cellular heterogeneity of transplanted PSC-CMs identifying a possible causal link between atrial and pacemaker like cardiomyocytes and arrhythmogenesis in PSC-CM treated subjects.

### Early activation of retinoic acid signalling during PSC-CM differentiation enriches for atrial and pacemaker-like subpopulations

To further explore the arrhythmogenic potential of atrial and pacemaker like cardiomyocytes, we developed a novel bioreactor differentiation protocol to enrich for these subpopulations. By modifying our standard protocol to activate the retinoic acid (RA) signalling pathway from day 2 to day 6, (Supplementary Fig. 1a and 7a) cardiac progenitors were directed to an atrial and pacemaker like fate (RA-PSC-CM)^39^. Expression of atrial and nodal genes such as *NPPA*, *MYL2a*, *SHOX2* and *HCN4* were upregulated as early as day 8 and significantly by day 15 in RA treated cultures, with converse suppression of ventricular markers such as *IRX4* and *MLC2v* (Supplementary Fig. 7d-f). This transcriptional pattern was confirmed with scRNA-seq analysis, in which RA-PSC-CMs showed a striking increase in atrial and pacemaker like cardiomyocytes and reduction of ventricular cardiomyocytes (Fig. 4a-b). Electrophysiological analysis confirmed the phenotypic difference between standard and RA-PSC-CMs cell preparations with RA-PSC-CMs exhibiting faster spontaneous firing rates, reduced action potential durations and lower sodium current densities (Fig. 4c-d), features all consistent with atrial and pacemaker rather than ventricular cardiomyocytes.

**Figure 4.**
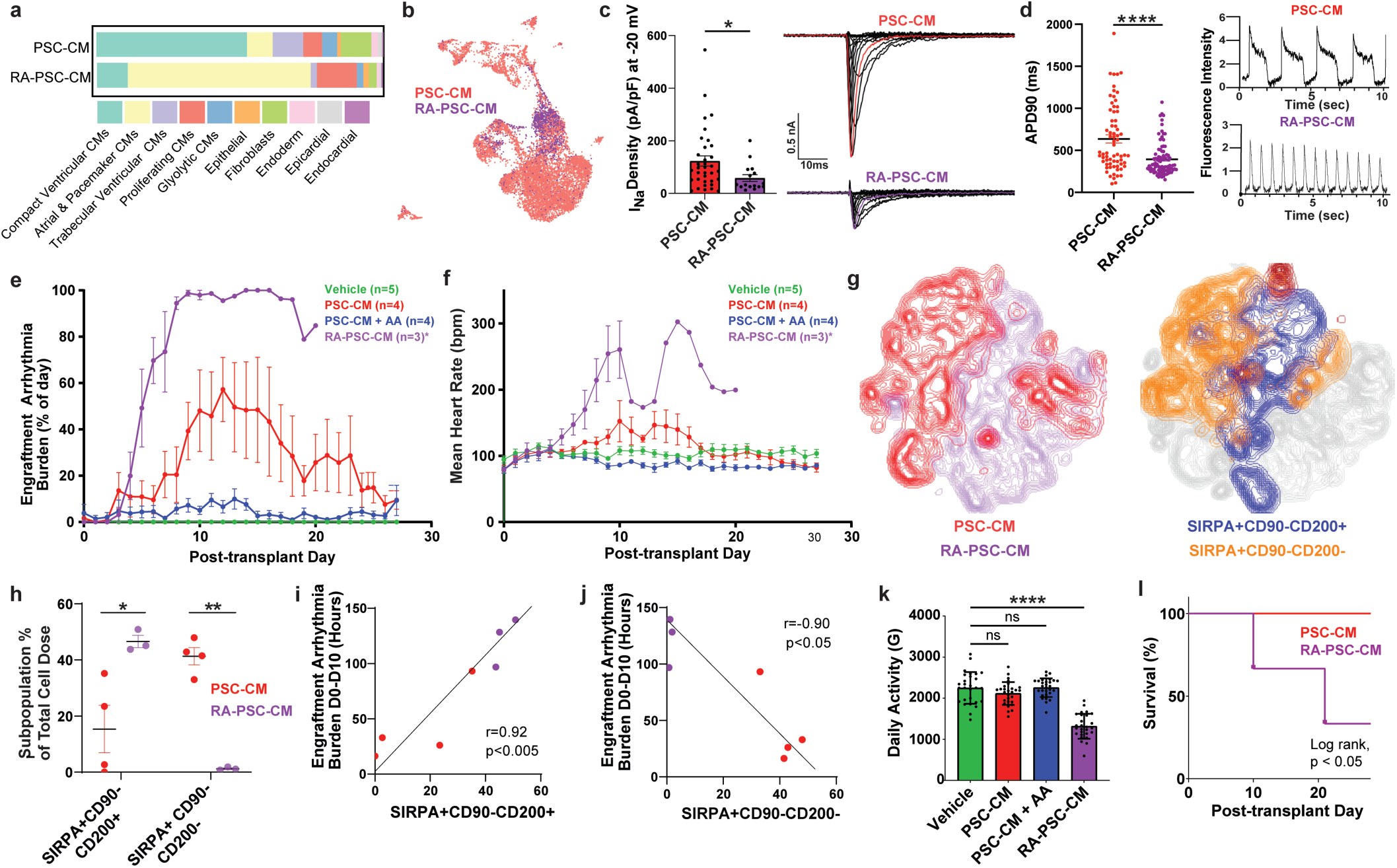
RA-PSC-CMs are enriched with atrial and pacemaker subpopulations and are highly arrhythmogenic. **(a)** Bar plot showing the contribution of different clusters to cells from representative PSC-CM and RA-PSC-CM cell doses. Proportionally greater contribution of compact ventricular cardiomyocytes in PSC-CM cell dose compared to greater proportion of atrial and pacemaker cardiomyocytes in RA-PSC-CM cell dose. **(b)** UMAP plot grouped by cells derived from either PSC-CM cell doses (red) or RA-PSC-CM cell doses (purple). RA-PSC-CM cell doses were predominantly made up of cluster 1 cells, atrial and pacemaker cardiomyocytes. **(c)** Analysis of sodium current densities in PSC-CMs and RA-PSC-CMs. Left - plot depicting maximum current densities recorded at -20mV in PSC-CMs (n = 35) and RA-PSC-CMs (n = 16). Significantly lower current densities in RA-PSC-CMs (* p = 0.02, Mann Whitney test). Right - Representative recordings of sodium current in PSC-CMs and RA-PSC-CMs made at different membrane potentials (inset: voltage protocol). **(d)** Left - Plot depicting action potential duration at 90% depolarisation (APD90) in PSC-CMs (n= 68) and RA-PSC-CMs (n=85). Significantly shorter APD90 in RA-PSC-CMs (**** p<0.0001, Mann Whitney test). Right - Representative appearance of action potential morphology and spontaneous beat rates from PSC-CMs and RA-PSC-CMs. **(e)** Pooled data expressing percentage of time spent in ventricular arrhythmia and **(f)** mean heart rate per day between groups (mean ± SEM). Substantially faster and more abundant arrhythmias in RA-PSC-CM group (*RA-PSC-CM n=3 only from day 0 - day 10, n=1 from day 11 - day 20. RA Pig #1 died on day 20, RA Pig #2 underwent catheter ablation on day 10 with data excluded thereafter, RA Pig #3 died on day 10). **(g)** High-parameter flow cytometric analysis comparing representative PSC-CM and RA- PSC-CM cell doses. Left: Concatenated tSNE plot showing distribution of cells from PSC-CM dose (red) and RA-PSC-CM dose (purple). Right: Concatenated tSNE plot showing distribution of SIRPA^+^CD90^-^CD200^+^ and SIRPA^+^CD90^-^CD200^-^ cardiomyocytes. Arrhythmogenic SIRPA^+^CD90^-^ CD200^+^ cardiomyocytes predominantly belong to RA-PSC-CM cell dose. Conversely non- arrhythmogenic SIRPA^+^CD90^-^CD200^-^ cardiomyocytes predominantly belong to PSC-CM cell dose. **(h)** Plot depicting proportion of SIRPA^+^CD90^-^CD200^+^ and SIRPA^+^CD90^-^CD200^-^ cardiomyocytes in each PSC-CM and RA-PSC-CM cell dose. Significantly increased proportion of arrhythmogenic SIRPA^+^CD90^-^CD200^+^ and reduced proportion of SIRPA^+^CD90^-^CD200^-^ cardiomyocytes in RA-PSC-CM doses (* p = 0.02, *** p = 0.0001, unpaired *t*-test, df = 5) **(i)** Correlation between total hours of arrhythmia from day 0 - day 10 and proportion of arrhythmogenic SIRPA^+^CD90^-^CD200^+^ cardiomyocytes or **(j)** non-arrhythmogenic SIRPA^+^CD90^-^CD200^-^ cardiomyocytes in PSC-CM (red) and RA-PSC-CM (purple) cell doses. Significant positive (r = 0.92, p = 0.003) and negative linear correlations (r = -0.90, p = 0.006). (k) Total daily activity between groups as measured in G-forces by accelerometers in implanted telemetry units. No significant difference in total activity between PSC- CM or PSC-CM + AA groups compared to vehicle, however RA-PSC-CM subjects had significantly lower daily activity (Vehicle vs PSC-CM: p = 0.24; Vehicle vs PSC-CM + AA: p > 0.99; Vehicle vs RA-PSC-CM: **** p < 0.0001; Ordinary one-way ANOVA with Dunnett’s multiple comparisons test) (l) Kaplan-Meier curve for overall survival showing RA-PSC-CM treatment is associated with reduced survival after 4 weeks (* p = 0.04, Log-rank test).

### RA-PSC-CM are highly arrhythmogenic after transplantation into infarcted myocardium

To confirm whether RA-PSC-CM transplantation would augment in vivo arrhythmia burden, transplantation studies were conducted in 3 further infarcted porcine subjects in the second phase of our large animal experiments. These additional animals all received an equivalent dose of 750 million RA- PSC-CMs 2 weeks following myocardial infarction. A striking increase in arrhythmia burden and rate was noted in all 3 animals, who experienced rapid and near continuous EA by day 8 post cell delivery (Fig. 4e-f). High parameter flow cytometric analysis of each cell dose showed a greater proportion of the suspected arrhythmogenic SIRPA^+^/CD90^-^/CD200^+^ subpopulation, and reduction in the non-arrhythmogenic SIRPA^+^/CD90^-^/CD200^-^ subpopulation (Fig. 4g-h). Importantly, these additional animals significantly strengthened the positive (r=0.92, p < 0.005) and negative (r=-0.90, p < 0.05) arrhythmia correlation of these surface marker signatures (Fig. 4i-j), confirming the potential utility for these signatures in arrhythmia prediction. The elevated arrhythmia rate and burden was less favourably tolerated by the animals, who exhibited significantly reduced activity levels as quantified by accelerometer data (Fig. 4k). Unfortunately, 2 of the 3 RA-PSC-CM recipients succumbed to either heart failure related or arrhythmic deaths prior to completing their experimental time course (Fig. 4l), with the third surviving only due to timely intervention with catheter ablation, described further below. Together, these data confirm the highly arrhythmogenic potential of atrial and pacemaker like cardiomyocytes, and identify SIRPA^+^/CD90^-^/CD200^+^ and SIRPA^+^/CD90^-^/CD200^-^ as novel surface marker signatures selecting for arrhythmogenic or non-arrhythmogenic cell preparations respectively.

### RA-PSC-CM grafts comprise of arrhythmogenic subpopulations, with reduced sarcomeric protein and intercalated disk organisation in comparison to PSC-CM grafts

The fate of engrafted cells was also probed in histological experiments. Human graft was identified by staining with antibodies against the human nuclear antigen Ku80 or by targeting the green fluorescence protein (GFP) within the gCaMP indicator of our cell line (Supplementary Fig. 8a). For detection of ventricular and atrial-like CMs within the graft, we stained sections with antibodies targeting cTnT, MLC2v and MLC2a. Interestingly, RA-PSC-CM graft comprised strikingly more MLC2a^+^ cardiomyocytes suggesting greater atrial cardiomyocyte engraftment (Fig. 5a, Supplementary Fig. 8b & 9a). In addition, these grafts also had a greater proportion of arrhythmogenic SIRPA^+^/CD200^+^ cardiomyocytes (Fig. 5b, Supplementary Fig. 9b).

**Figure 5.**
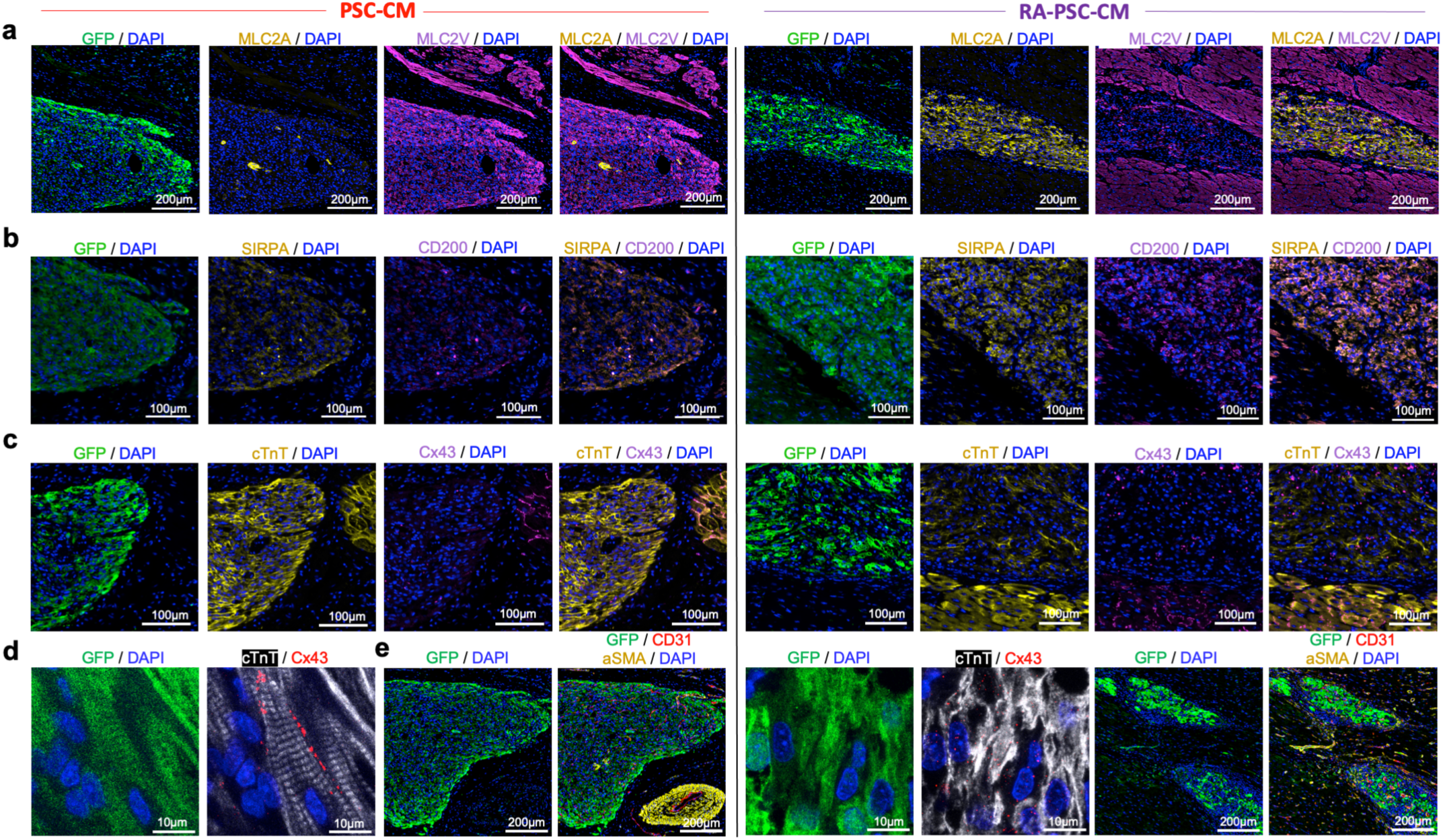
RA-PSC-CM grafts are enriched for atrial myocyte markers and show reduced sarcomeric protein and connexin 43 organisation in comparison to PSC-CM grafts. **(a)** Representative immunofluorescence images from PSC-CM and RA-PSC-CM treated hearts showing PSC-CM grafts are composed almost entirely of MLC2v+ myocytes, in contrast to RA-PSC-CM grafts, which predominantly contain MLC2a+ myocytes. **(b)** Representative immunofluorescence images from PSC-CM and RA-PSC-CM treated hearts, showing increased abundance of arrhythmogenic SIRPA+/CD200+ myocytes in RA-PSC-CM grafts. **(c)** Low magnification immunofluorescent images of cardiac troponin T (cTnT) and connexin 43 (Cx43) staining in PSC-CM and RA-PSC-CM grafts, showing increased expression of cTnT in PSC-CM grafts compared to RA- PSC-CM. In both graft types, Cx43 expression is less abundant than in the surrounding pig myocardium. **(d)** High magnification confocal images of the previous grafts showing highly organised and aligned sarcomeres with appropriate localisation of Cx43 to the intercalated disks in PSC-CM graft. In stark contrast, RA-PSC-CM graft has disorganised cTnT expression, with sporadic expression of Cx43. **(e)** Immunofluorescent staining for CD31 and alpha smooth muscle actin shows an abundance of neovessels in both PSC-CM and RA-PSC-CM grafts.

To visualise the organisation of sarcomeres and the formation of gap junctions between cardiomyocytes, we stained grafts with antibodies against cTnT and connexin 43 (Cx43) (Fig. 5c & Supplementary Fig. 9c). High magnification confocal images show organised sarcomeres in standard PSC-CM grafts, with appropriate localisation of Cx43 to the intercalated disks (Fig. 5d & Supplementary Fig. 8c). In stark contrast, RA-PSC-CM grafts had disorganised cTnT expression, with sporadic and lateralised Cx43 expression patterns compared to standard PSC-CM grafts.

We also stained tissue with antibodies against CD31 and alpha smooth muscle actin to identify new vessel growth within the grafts (Fig. 5e, Supplementary Fig. 9d). Both standard PSC-CM and RA-PSC-CM grafts contained abundant micro-vessels (capillaries and arterioles), important in supporting long-term graft survival.

### Catheter ablation is a feasible alternative treatment strategy for PSC-CM related engraftment arrhythmias

Despite our demonstration of amiodarone-ivabradine supressing EA burden, a fall-back strategy in the case of aggressive or refractory EAs is imperative for safe PSC-CM clinical translation. Thus, in the third and final phase of our large animal studies, we sought to determine the feasibility and efficacy of catheter ablation (CA) as an alternative EA treatment strategy. A further 2 pigs underwent PSC-CM delivery following myocardial infarction, with the intention of proceeding to CA 2 weeks post cell injection. Only one of these subjects displayed sufficient EA to facilitate electroanatomic mapping and ablation, with inadequate burden in the second (Supplementary Fig. 10a). ScRNA-seq for the latter showed minimal contribution of atrial and pacemaker-like cardiomyocytes in the input cell dose with flow cytometry showing a high percentage of non-arrhythmogenic SIRPA^+^/CD90^-^/CD200^-^ cardiomyocytes. Together these findings account for the low arrhythmia burden in this pig (Supplementary Fig. 10b-c). In the first animal, EA burden was trending upward by day 13, at which point electroanatomic mapping and CA was performed. This localised EA origin to the inferolateral apex, and this region was targeted with a series of ablations terminating EA and restoring sinus rhythm (Figure 6a-b). EA did not recur intraoperatively despite a period of monitoring and aggressive arrhythmia induction strategies (programmed electrical stimulation, isoprenaline infusion, burst pacing). Telemetry analysis from the subsequent 2 weeks showed a marked drop in EA burden (Fig. 6c) with only isolated ventricular ectopic beats rather than sustained arrhythmias noted.

**Figure 6.**
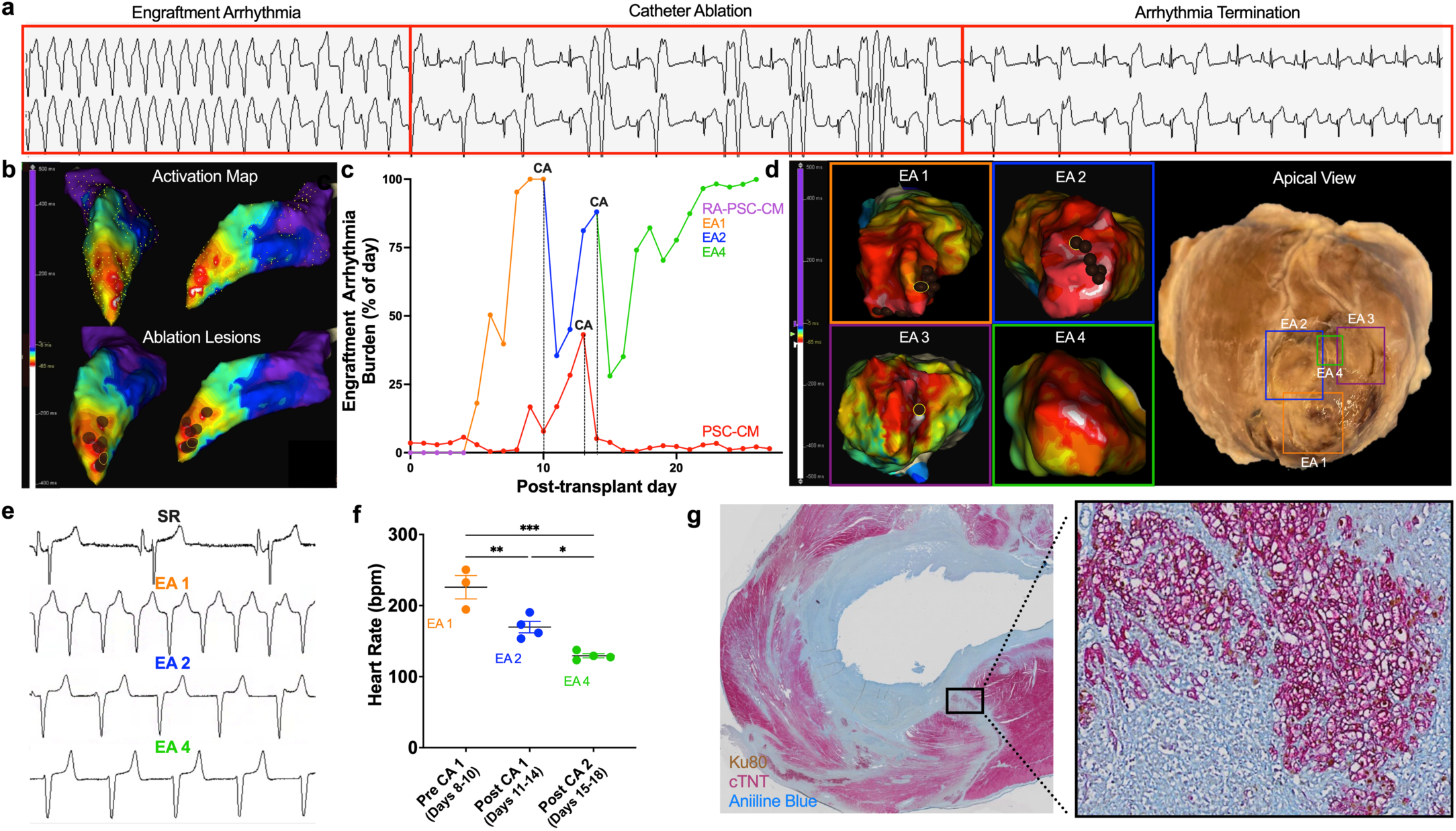
Catheter ablation is a feasible and effective EA treatment strategy. **(a)** Representative rhythm strip showing termination of EA during CA. **(b)** Electroanatomic maps from representative CA treated subject. Top - Activation map of EA showing anatomic origin of arrhythmia (early activation - white, late activation - purple). Bottom - Activation map overlaid with ablation lesions (brown circles) which were delivered at sites of earliest activation resulting in termination of arrhythmia. **(c)** Plot depicting percentage of day spent in ventricular arrhythmia in subjects treated with catheter ablation. Two ablation procedures required for RA-PSC-CM subject and one procedure required for PSC-CM subject. **(d)** Anatomic correlation between endocardial activation maps and extracted heart in RA-PSC-CM subject. Left - Representative activation maps in which 4 unique EAs were encountered. EAs 1-3 were terminated during CAs 1 and 2 with final ablation lesions highlighted (brown circle with yellow outline). EA4 was persistent after CA2 and mapped at terminal EPS. Right - Apical view of extracted heart following euthanasia and formalin fixation. Ablation lesions (EAs 1-3) and island of surviving cell graft (EA4) outlined with excellent anatomic correlation to preceding activation maps. **(e)** Representative rhythm strips from RA-PSC-CM subject of sinus rhythm, EA 1, EA 2, and EA 4. **(f)** Plot depicting daily heart rate of EA 1, 2 and 4 in RA-PSC-CM subject (mean ± SEM). Each new EA was of a significantly slower heart rate than the previously ablated EAs (EA 1 vs EA 2 - ** p = 0.01, EA 1 vs EA 4 - *** p = 0.0003, EA 2 vs EA 4 - * p = 0.04; Ordinary one-way ANOVA with Sidak’s multiple comparisons test). **(g)** Whole-mount cross-section of *RA-PSC-CM + CA* subject. Engrafted human cardiomyocytes expressed cardiac troponin T (red) and the human-specific anti-nuclear antigen Ku80 (brown). Scarred myocardium, either from percutaneous myocardial infarction or catheter ablation, was identified with aniline blue counterstaining. A small region of surviving human cardiomyocyte graft responsible for residual arrhythmias was identified (boxed region, zoomed on right).

After this successful proof-of-concept experiment, we sought to assess the efficacy of CA in pigs treated with the highly arrhythmogenic RA-PSC-CMs. Only 1 of the 3 pigs treated with RA-PSC-CM underwent CA, with the remaining 2 animals dying before planned ablation procedures could take place. In the ablated animal, CA was a lifesaving intervention facilitating survival through to the terminal time-point. However, two procedures were required with a greater number of ablation lesions and higher radiofrequency doses required for arrhythmia termination compared to the standard PSC-CM pig (Supp. Table 6). In addition, EA recurred within 12 hours after each procedure despite apparent initial procedural success (Fig. 6c). Interestingly, recurrent EA was always found to originate from a new site, correlating with a separate cell injection location. In total, 4 different origins of EAs were identified, 3 of which (EAs 1-3) were successfully ablated. The final arrhythmia (EA4) was mapped at the terminal procedure but not ablated due to the pre-determined endpoint being reached. Examination of the heart following tissue harvest demonstrated excellent anatomic correlation with preceding endocardial activation maps with clearly visible transmural ablation lesions identifying EA1- 3 along with an island of surviving cell graft identifying EA4 (Fig. 6d). Of interest, despite recurrence following each ablation, the new arrhythmias were of lower heart rates (Fig. 6e-f) and more favourably tolerated by the animal on clinical assessment. Histological analysis confirmed successful disruption of engrafted regions with each ablation, with intact, surviving graft responsible for the residual arrhythmia (Fig. 6g).

Taken together, these results indicate that CA is a feasible therapeutic strategy for EA, however PSC-CM grafts may demonstrate hierarchical pacemaker-like attributes with ectopic EA activity falling back to alternative graft sites once the dominant graft has been ablated. Treatment success, particularly with highly arrhythmogenic cell doses, may require complete ablation of all engrafted regions if arrhythmogenic cell populations are not removed prior to transplantation.

## Discussion

The prospect of significant cardiac remuscularisation with PSC-CMs offers renewed hope to heart failure patients^14–18^. Clinical trials are now being planned or in progress globally. However, pre-clinical data informing on specific key areas could expedite successful clinical translation. In this regard several studies using clinically relevant large animal models have shown that EAs are a predictable complication following intramyocardial PSC-CM delivery^8–12^. Here we show that PSC-CM related EAs can be suppressed and potentially abolished with clinically available pharmacological and procedural therapeutics. We further identify cellular characteristics of arrhythmogenic PSC-CMs through robust phenotyping of input and engrafted cells. These data can inform cell production strategies to provide safe and effective PSC-CMs for future clinical trials.

We show that the combination of ivabradine and amiodarone is extremely effective in supressing graft automaticity as well as reducing EA rate and burden. This result was independently reported^12^ whilst the current study was in progress. Ivabradine was selected given its specific inhibition of the I_f_ current responsible for pacemaker automaticity^44–46^ a characteristic we hypothesised may be effective in supressing automatic EAs. Conversely, amiodarone was selected given it is an established anti- arrhythmic agent which may be empirically administered in future PSC-CM clinical trials. It has a broad mechanism of action, antagonising K^+^, Na^+^ and Ca^2+^ channels along with β-adrenergic receptors^47^. The combination of these drugs effectively supressed but did not eradicate EAs, necessitating assessment of alternate treatment strategies particularly in the case of highly arrhythmogenic cell doses. Therefore, we went on to show that catheter ablation is a feasible strategy to treat and abolish EA. This proof-of-concept result has important practical implications as the first patients are being treated with PSC-CMs. Interestingly, our ablation experiments also yielded important EA mechanistic insights, demonstrating that pacemaker activity can fall back to grafts of lower intrinsic rates once the dominant graft has been ablated. A caveat to CA therapy for EA is that ablating multiple graft regions risks loss of the contractile benefits exerted by PSC-CM transplantation. As such, we propose CA only be pursued in the case of aggressive and pharmacologically refractory EAs and more prudently, that generation of less arrhythmogenic PSC-CM cell products be investigated. To this end, we also sought to gain mechanistic insights into the cellular compositions which may contribute to EAs.

Substantial variability of EA burden has been reported within and between PSC-CM large animal studies^8–12^. Though this may be reflective of several factors such as differences in animal species or cell delivery technique, it is likely to be predominantly determined by the molecular make-up of the cellular product^19^. Reductions in arrhythmia burden have been noted after a period of *in vivo* graft maturation^8, 9^, suggesting PSC-CM immaturity may be a key determinant of arrhythmogenesis^48–51^. However, not all PSC-CM recipients achieve ‘electrical maturation’^12^, implying immaturity may not be the only important cellular characteristic. Current PSC-CM differentiation protocols are known to generate heterogeneous cell populations containing a mix of ventricular, atrial and pacemaker like cardiomyocytes^22, 23^. For the first time, we outline the importance of this heterogeneity in arrhythmogenesis, identifying atrial and pacemaker-like cells as culprit subpopulations along with describing unique surface marker signatures predictive of arrhythmogenicity.

Cells enriched for atrial and pacemaker-like cardiomyocytes were generated through addition of retinoic acid to our bioreactor differentiation protocol^52, 53^ and shown to be highly arrhythmogenic, supporting a causal link between these subpopulations and EA. We show that the enhanced automaticity of these subpopulations *in vitro* directly translates to more abundant and rapid EAs *in vivo.* Furthermore, by robustly phenotyping input cell doses and correlating these to resultant arrhythmia burden, we identify SIRPA^+^/CD90^+^/CD200^+^ and SIRPA^+^/CD90^-^ /CD200^-^ cells as arrhythmogenic and non-arrhythmogenic cardiomyocytes respectively. In doing so, we not only provide a simple quality control tool for assessing the arrhythmogenic potential of cell doses, but also an avenue for removing arrhythmogenic cells by cell sorting prior to transplantation. Interestingly, our scRNA-seq data shows these arrhythmogenic SIRPA^+^/CD90^+^/CD200^+^ cardiomyocytes have a transcriptomic signature consistent with atrial and pacemaker subpopulations, further supporting the notion that these cell types are important in arrhythmogenesis. There is currently substantial interest in generating chamber-specific cardiomyocytes for various therapeutic or drug discovery applications^21, 24, 39, 52, 54–58^. Our data further endorses the pursuit of transplanting purified ventricular cardiomyocytes, devoid of atrial and pacemaker-like cells for the purposes of therapeutic cardiac remuscularisation.

Finally, although not our primary intention and despite small sample sizes, we were able to demonstrate left ventricular functional improvement in all cell recipients but particularly those in those which EAs had been ameliorated with pharmacotherapy. PSC-CM grafts elicit direct contractile force^59^ with our data suggesting that this beneficial effect can be further enhanced once arrhythmogenicity is addressed. Interestingly, a trend to improvement was also noted in the ungrafted right ventricle, suggesting immunomodulatory paracrine influences may also be a factor ^60^.

There are several limitations to this study. Firstly, because the primary focus was to study EAs rather than show functional improvement, both infarct and sample sizes were small. Secondly, as in most other large animal PSC-CM pre-clinical studies^8–11^ transplantation experiments were conducted in the sub- acute phase post-MI, despite planned clinical trials likely to be conducted in patients with chronic ischaemic heart failure ^18, 20^. Thirdly, only a single, embryonic stem cell line was assessed, with further work using induced PSC-CMs (iPSC-CMs) produced under good manufacturing practice an active pursuit of our group.

In conclusion, repopulation of infarcted myocardium with functional, force generating cardiomyocytes is an exciting therapeutic prospect positioning PSC-CMs as a leading candidate for cardiac regeneration. Though associated with arrhythmogenesis, here we deepen our mechanistic understanding of this predictable complication, informing that it is likely addressable through modifications to cardiomyocyte production protocols. Additionally, we show that PSC-CM engraftment arrhythmias can be suppressed with clinically available pharmacologic and procedural anti-arrhythmic strategies, an important safety consideration given several impending clinical trials.

## Acknowledgements

We would like to acknowledge all veterinary staff, animal support staff and research technical staff involved in this project. In particular we acknowledge Dr Rafael Dye, Dr Laurencie Brunel, Min Siok Teoh, Tuan Quang & Anh Vo.

We would also like to acknowledge Dr. Anton Shpak and Dr Satya Arjunan for the software code development of the KIC data analysis software (KIC DAT).

---

**Supplementary Figure 1.**
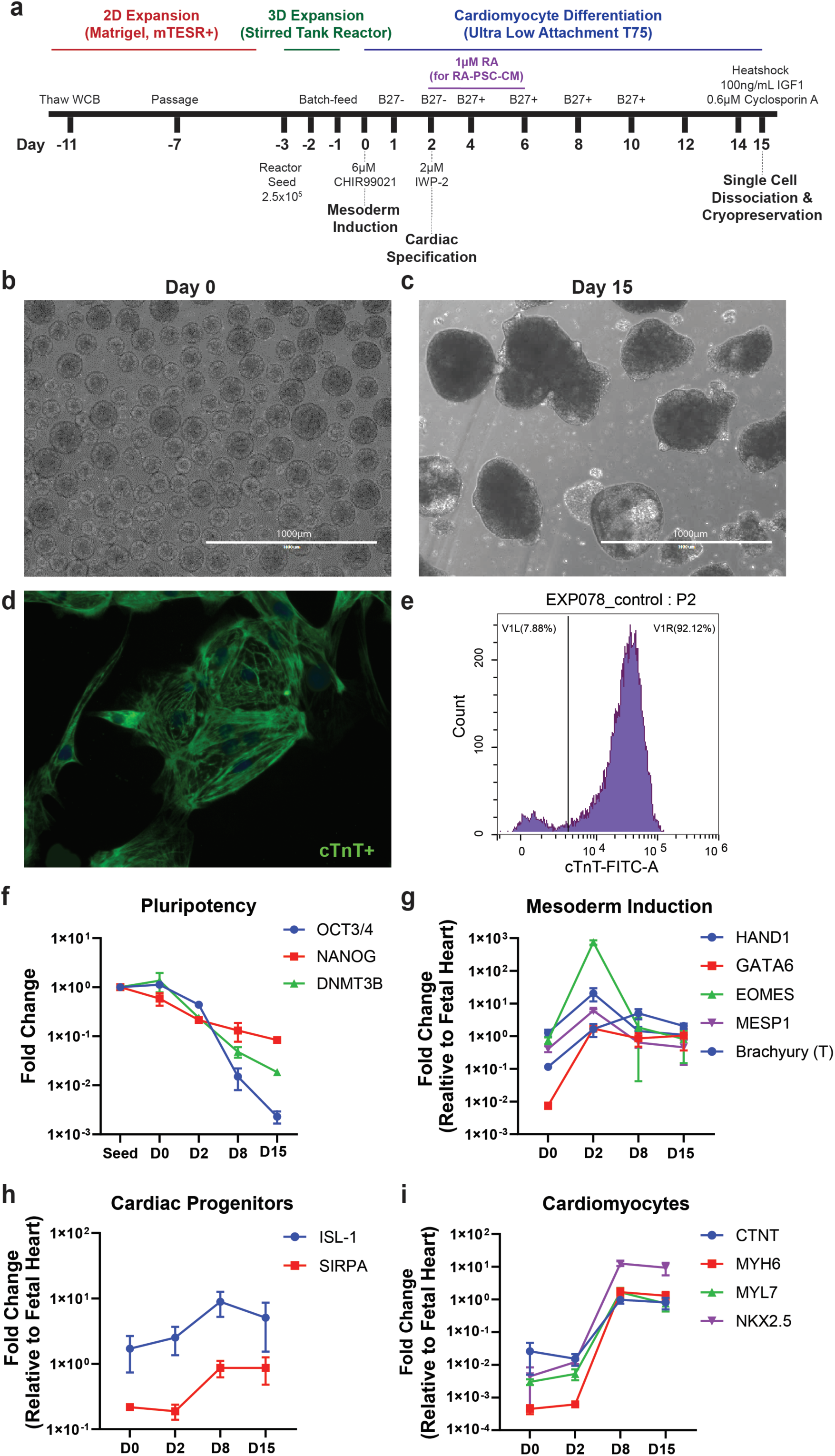
Stirred tank bioreactor production of PSC-CMs for porcine transplantation experiments. **(a)** Schematic depicting PSC-CM stirred tank reactor production protocol. **(b)** Appearance of cells at day 0 and **(c)** day 15 of differentiation protocol. **(d)** Immunostaining for cardiac troponin T demonstrating evidence of cytoskeletal formation in PSC-CMs. **(e)** Representative flow cytometric evaluation of PSC-CMs quantifying cardiac troponin T expression. **(f)** qPCR data showing expression profile of pluripotency, **(g)** mesodermal, **(h)** cardiac progenitor, and **(i)** cardiomyocyte markers over time course of differentiation protocol.

**Supplementary Figure 2.**
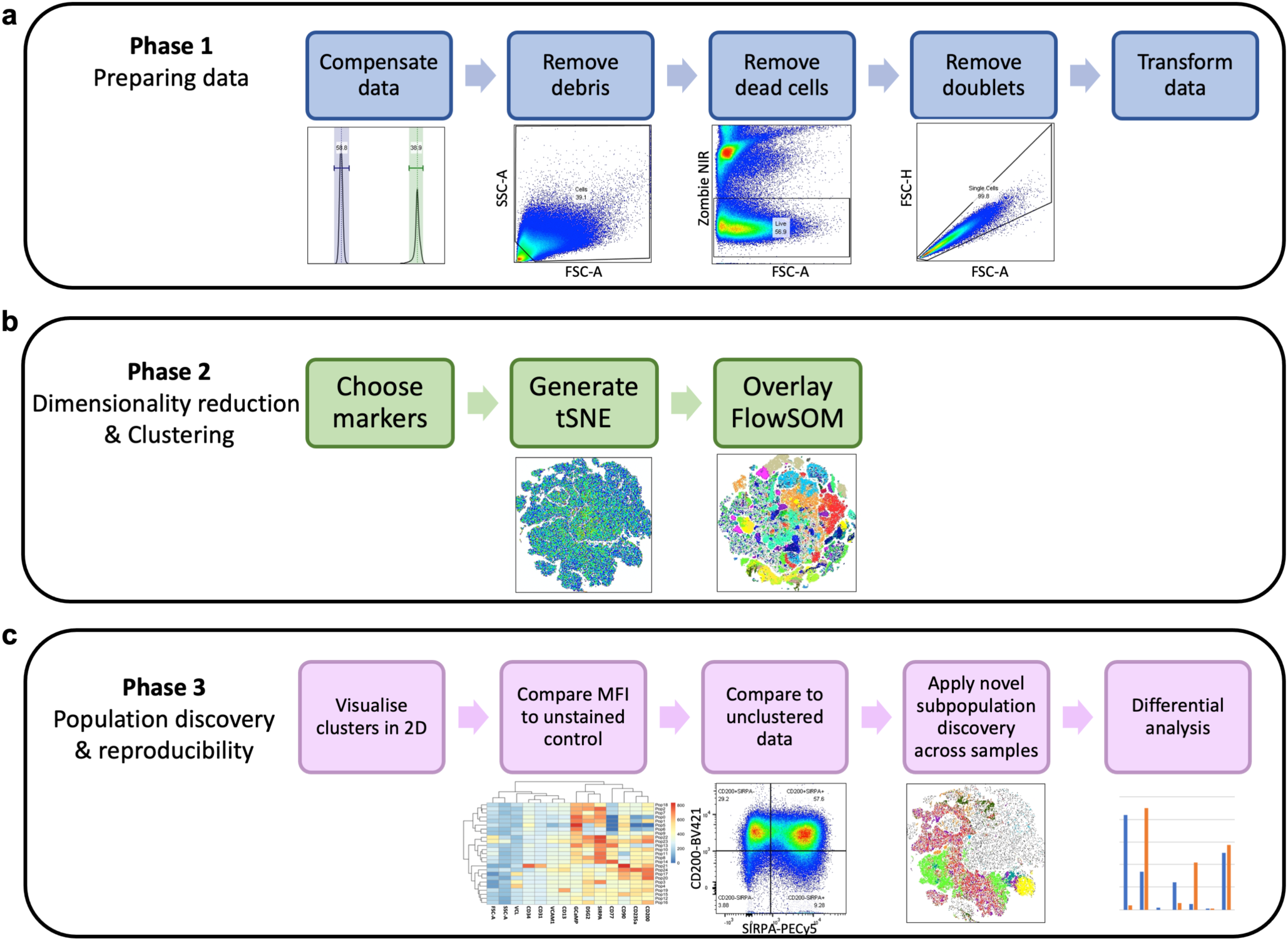
High parameter flow cytometry analysis workflow. **(a)** Steps involved in preparing data, **(b)** generating tSNE plot and overlaying with FlowSOM clusters, and **(c)** identifying and quantifying subpopulations.

**Supplementary Figure 3.**
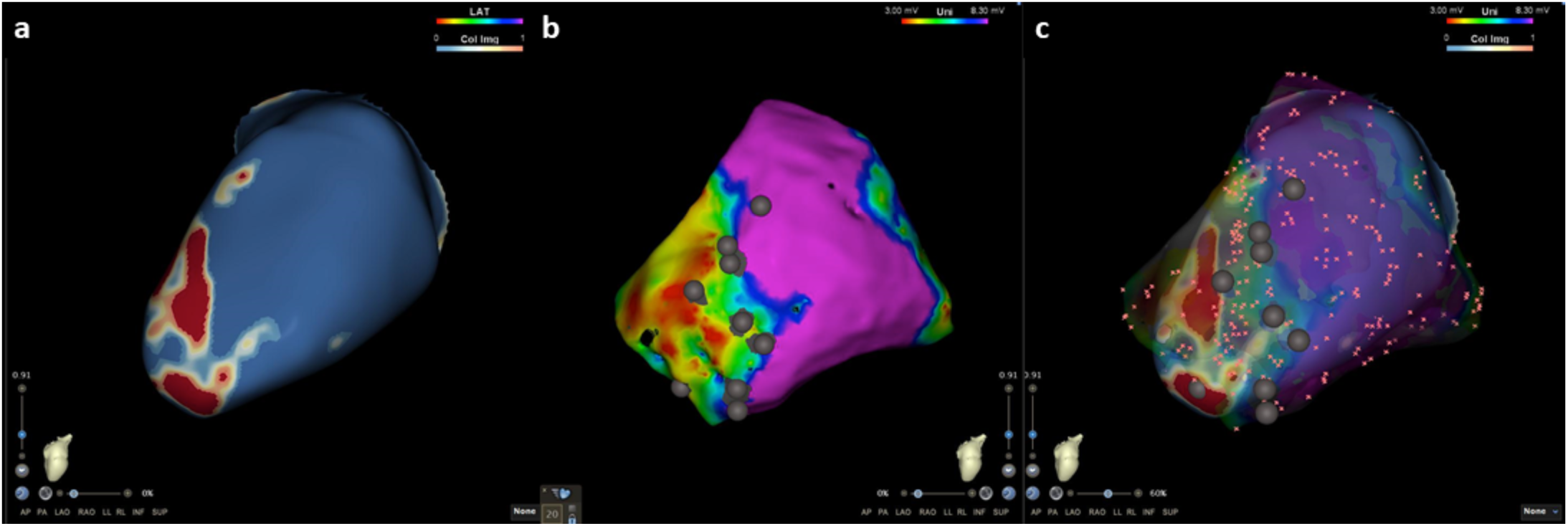
Merging of reconstructed CMR data and epicardial voltage map for localisation and annotation of cell injection sites. **(a)** Representative ADAS-3D map of left ventricle created through reconstruction of late gadolinium enhanced CMR data. Blue - healthy myocardium, Red - myocardial scar. **(b)** Epicardial map created by electroanatomically mapping left ventricle during thoracotomy procedure. Cell injections targeted to infarct and border zones and denoted by grey circles. Pink - healthy myocardium, blue/green - border zones, yellow/red - myocardial scar. **(c)** Merging of ADAS-3D map with epicardial voltage map performed using CartoMerge software providing accurate anatomic representation of left ventricle and cell injection sites.

**Supplementary Figure 4.**
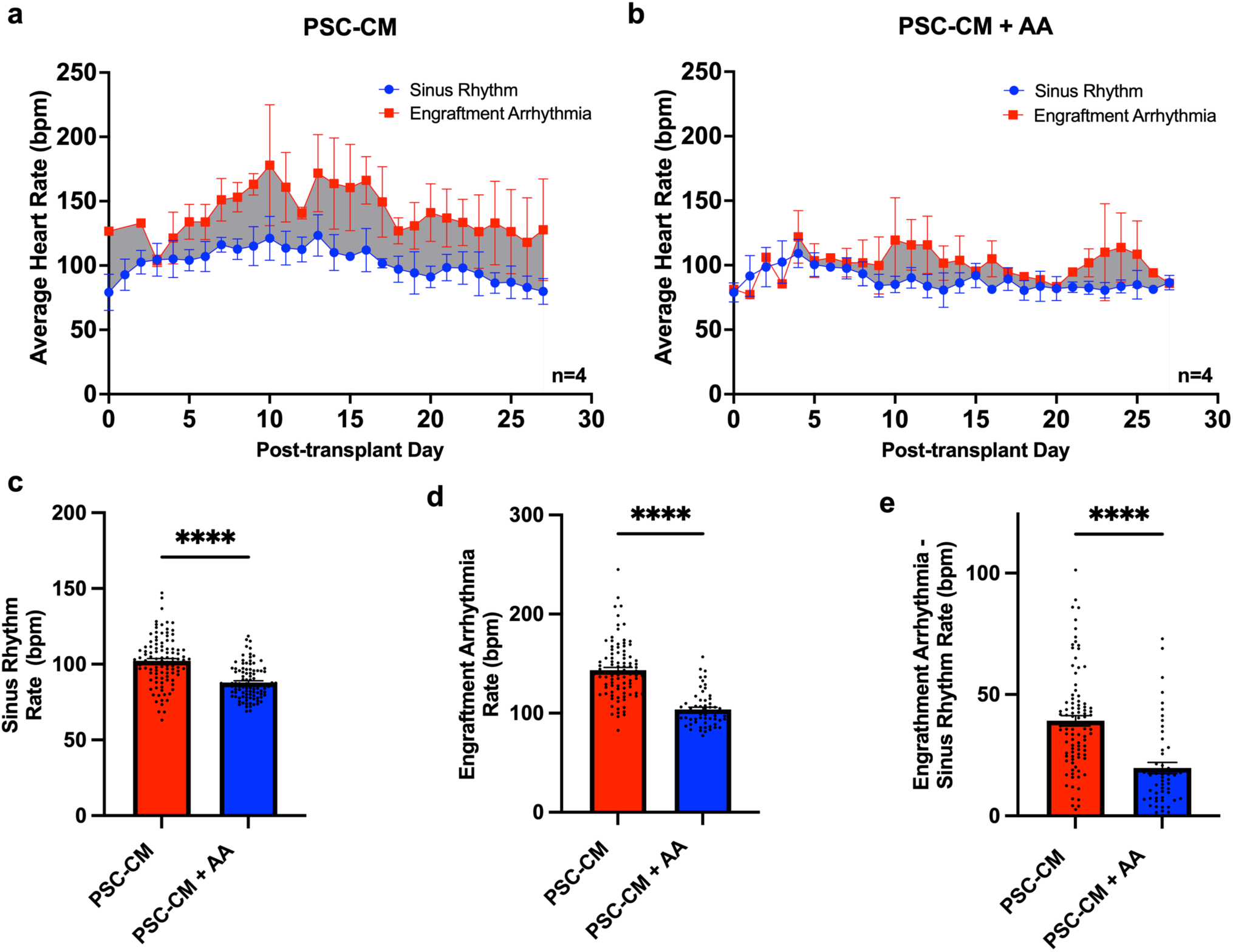
Anti-arrhythmic therapy reduces the average heart rate of sinus rhythm and engraftment arrhythmias in cell recipients. **(a)** Daily average heart rate of either sinus rhythm (blue) or engraftment arrhythmia (red) in PSC-CM recipients and **(b)** PSC-CM + AA recipients. **(c)** Significant reduction in rate of sinus rhythm, **(d)** engraftment arrhythmia, or **(e)** difference between engraftment arrhythmia and sinus rhythm in PSC-CM + AA versus PSC-CM subjects (****p<0.0001, Mann Whitney test).

**Supplementary Figure 5.**
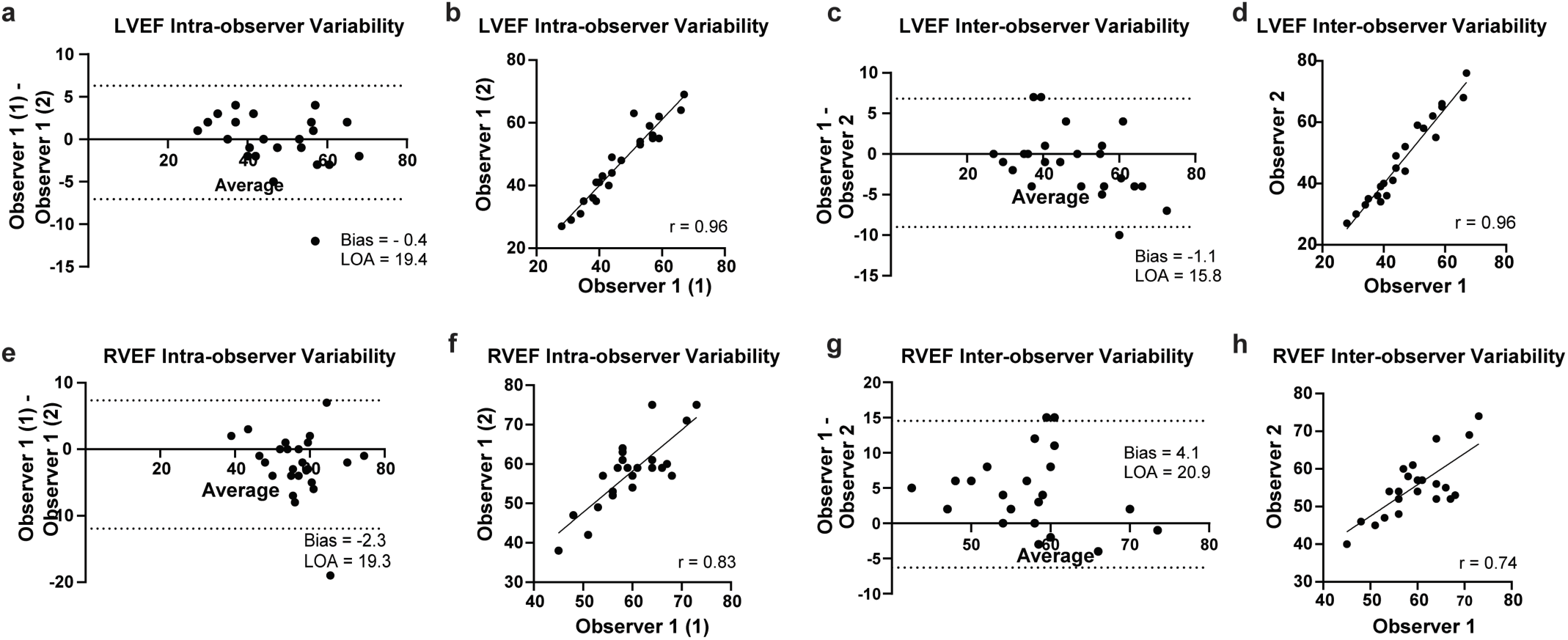
Reproducibility of CMR analysis. **(a)** Intra-observer variability of LVEF analysis depicted by Bland-Altman analysis and **(b)** correlation scatter plot. **(c)** Inter-observer variability of LVEF analysis depicted by Bland-Altman analysis and **(d)** correlation scatter plot. **(e)** Intra-observer variability of RVEF analysis depicted by Bland-Altman analysis and **(f)** correlation scatter plot. **(g)** Inter-observer variability of RVEF analysis depicted by Bland-Altman analysis and **(h)** correlation scatter plot.

**Supplementary Figure 6.**
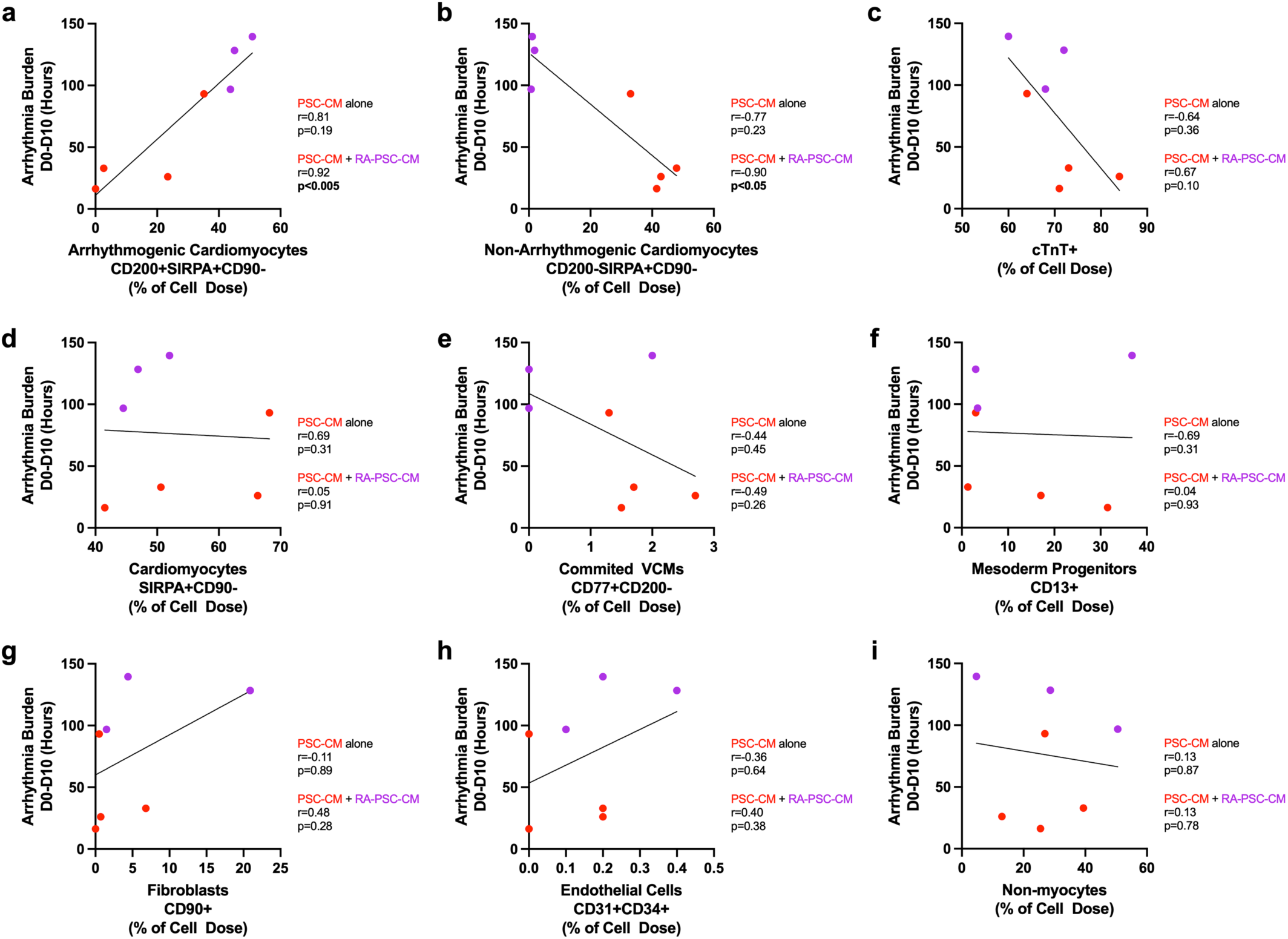
Correlation scatter plots demonstrating strength of linear association between flow cytometry derived subpopulation quantification and resultant arrhythmia burden for each PSC-CM (red) and RA-PSC-CM (purple) cell dose. **(a)** Arrhythmogenic cardiomyocytes, SIRPA^+^CD90^-^CD200^+^ **(b)** Non-arrhythmogenic cardiomyocytes, SIRPA^+^CD90^-^CD200^-^ **(c)** Cardiac troponin T positive cells **(d)** Cardiomyocytes, SIRPA^+^CD90^-^ **(e)** Committed ventricular cardiomyocytes, CD77^+^CD200^-^ **(f)** Mesodermal progenitors, CD13^+^ **(g)** Fibroblasts, CD90^+^ **(h)** Endothelial cells, CD31^+^CD34^+^ **(i)** Non-myocytes

**Supplementary Figure 7.**
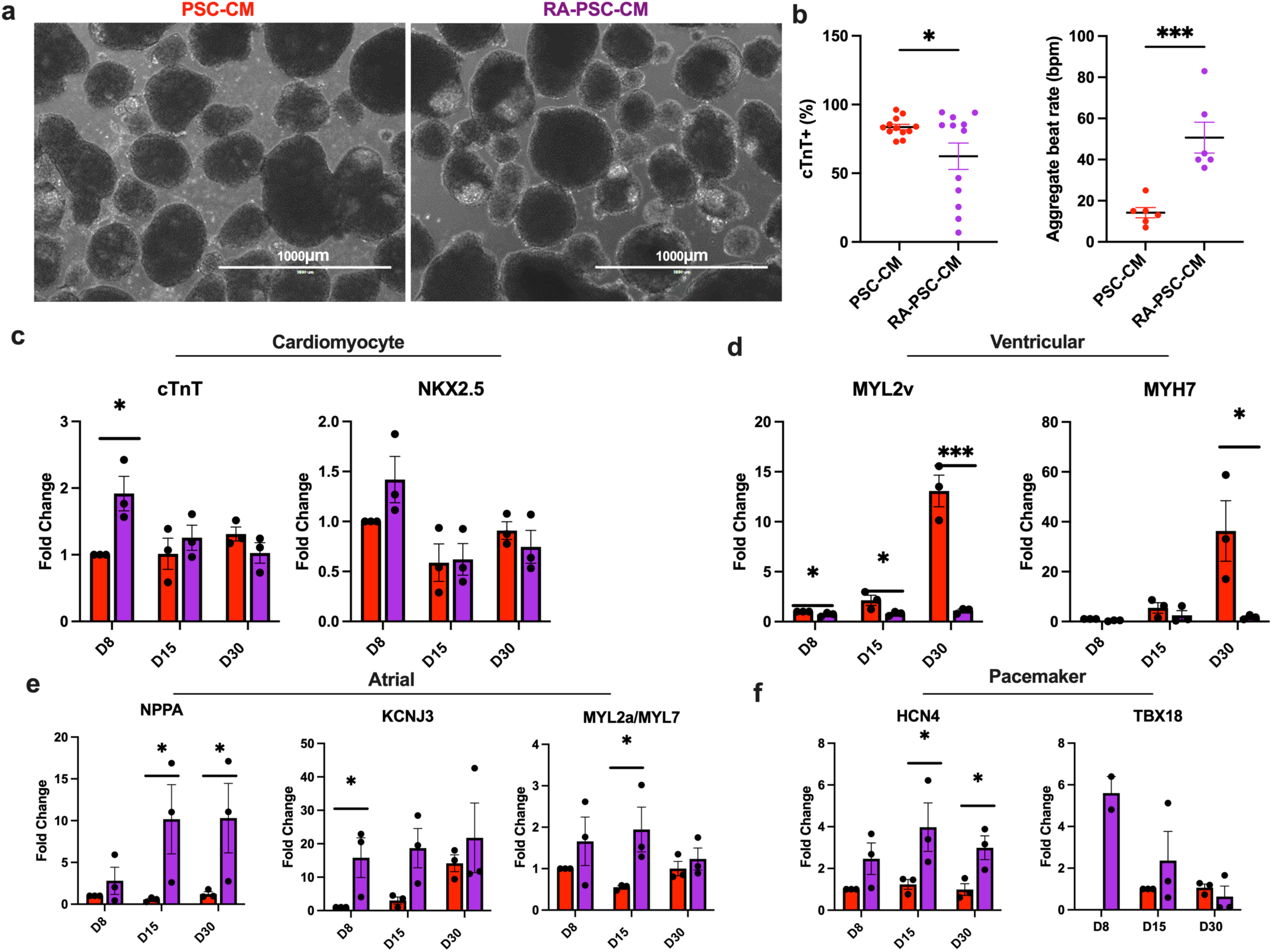
Stirred tank bioreactor generation and characterisation of RA-PSC-CMs. **(a)** Appearance of PSC-CM and RA-PSC-CMs at day 15 of bioreactor differentiation protocol. **(b)** Cardiac troponin T expression and **(c)** aggregate beat rate comparison between PSC-CMs and RA- PSC-CMs (*p<0.05, ***p<0.001, unpaired T test). **(c)** qPCR quantified expression of general cardiomyocyte, **(d)** ventricular, **(e)** atrial, and **(f)** pacemaker cardiomyocyte markers (* p<0.05, ***p<0.001, unpaired T test).

**Supplementary Figure 8.**
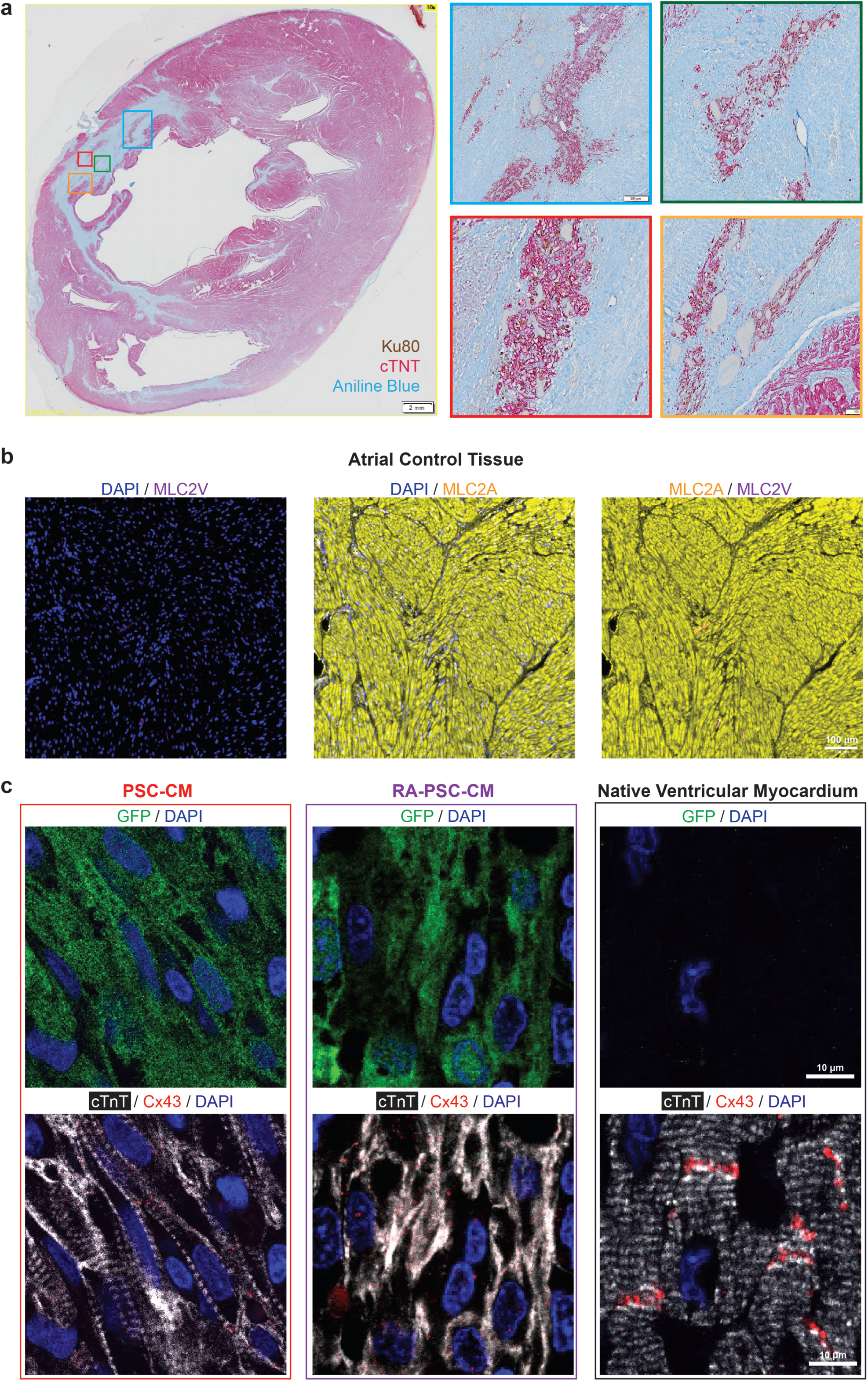
PSC-CM and RA-PSC-CM graft histology. **(a)** Representative whole-mount cross-section through PSC-CM recipient heart 28 days post transplantation. Engrafted human cardiomyocytes expressed cardiac troponin T (red) and the human-specific anti-nuclear antigen Ku80 (brown). Infarct was identified with aniline blue counterstaining. **(b)** MLC2A and MLC2V antibodies were optimised using control porcine atrial tissue (MLC2A+/MLC2V-). **(c)** Confocal microscopy of PSC-CM graft (left), RA-PSC-CM graft (middle) and native porcine ventricular myocardium (right). Top panels identify engrafted GFP+ human cells. Bottom panels show expression of cardiac troponin T and connexin 43 in engrafted regions compared with native ventricular myocardium. Standard PSC-CM graft showed organised and aligned sarcomeric structures with appropriate localisation of Cx43 to the intercalated disks. In stark contrast, RA-PSC-CM grafts had disorganised cTnT expression, with reduced Cx43 compared to standard PSC-CM grafts, and lateralised expression pattern.

**Supplementary Figure 9.**
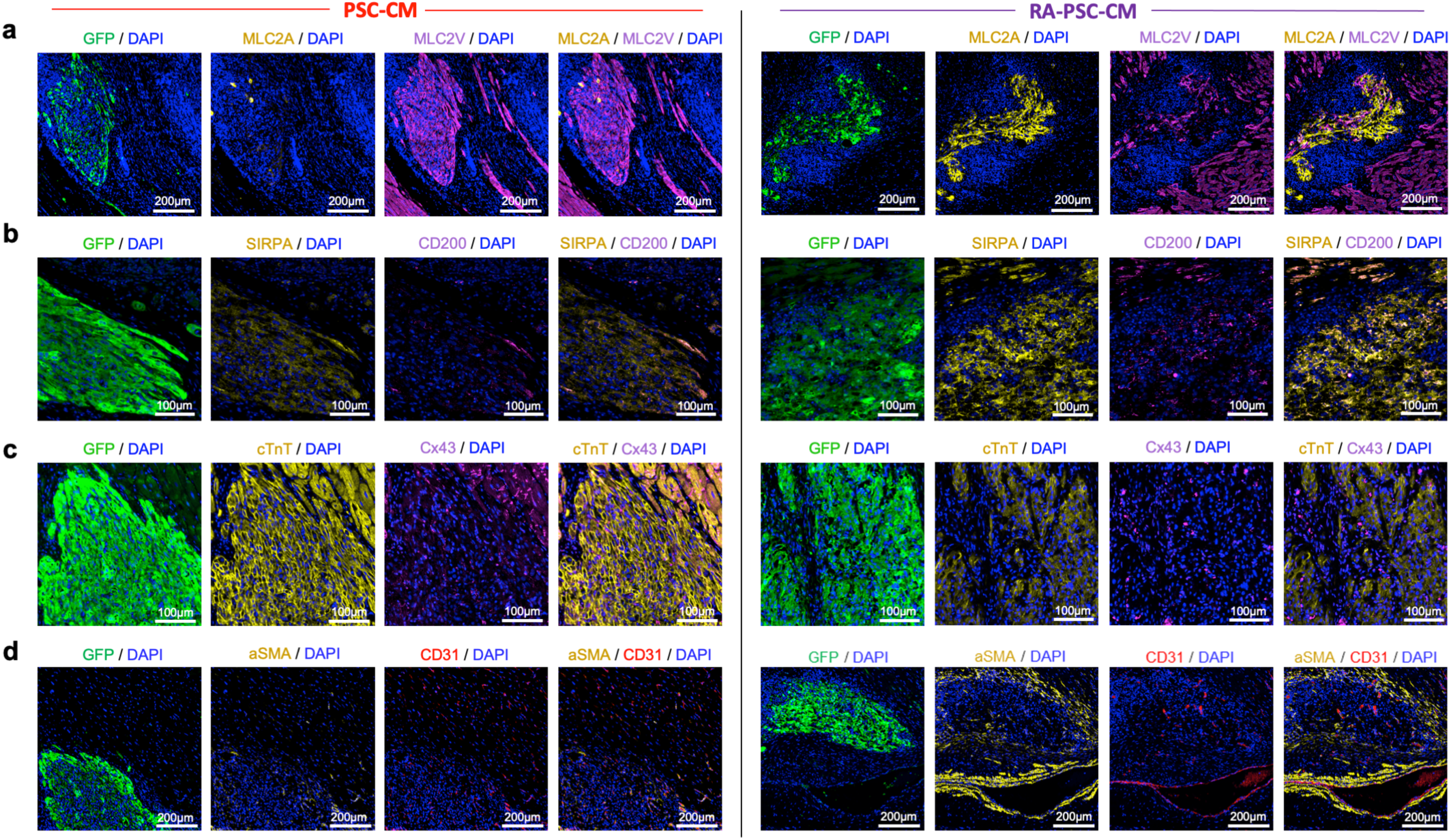
Additional PSC-CM and RA-PSC-CM graft histology. **(a)** Representative immunofluorescence images from additional PSC-CM and RA-PSC-CM treated hearts, showing PSC-CM grafts are composed almost entirely of MLC2v+ myocytes, in contrast to RA-PSC-CM grafts, which predominantly contain MLC2a+ myocytes. **(b)** Representative immunofluorescence images from PSC-CM and RA-PSC-CM treated hearts, showing increased abundance of arrhythmogenic SIRPA+/CD200+ myocytes in RA-PSC-CM grafts. **(c)** Low magnification immunofluorescent images of cardiac troponin T (cTnT) and connexin 43 (Cx43) staining in PSC-CM and RA-PSC-CM grafts, showing increased expression of cTnT in PSC-CM grafts compared to RA-PSC-CM. In both graft types, Cx43 expression is less abundant than in the surrounding pig myocardium. **(d)** Immunofluorescent staining for CD31 and alpha smooth muscle actin shows an abundance of neovessels in both PSC-CM and RA-PSC-CM grafts.

**Supplementary Figure 10.**
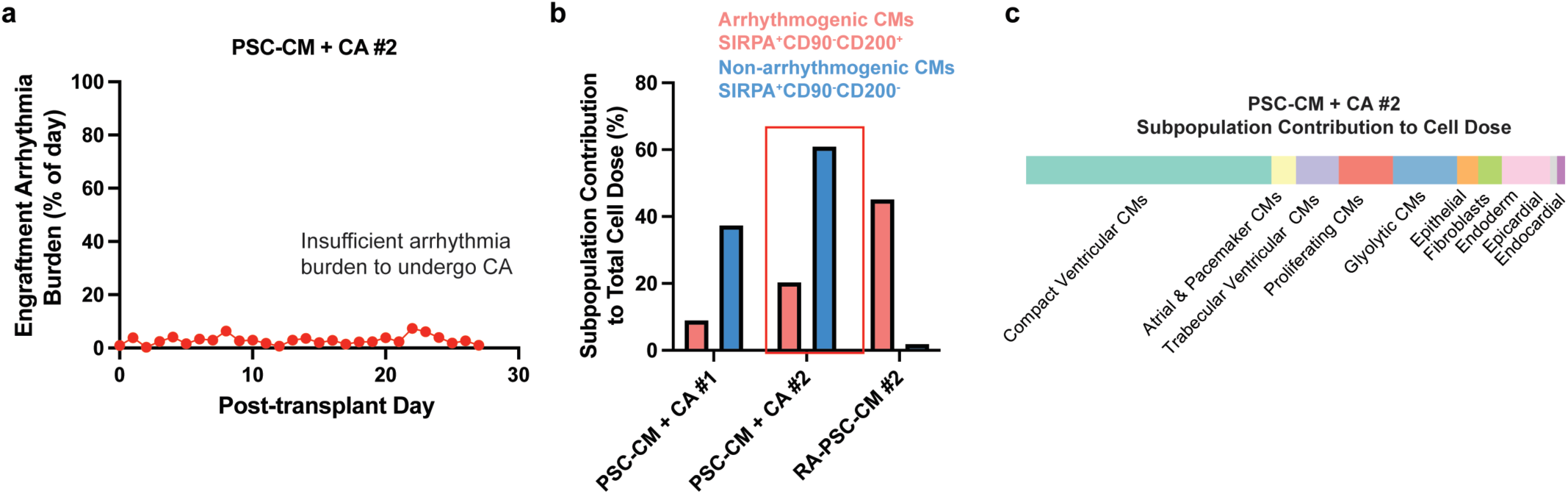
Flow cytometric and scRNA-seq profiling of PSC-CM cell dose with insufficient resultant engraftment arrhythmia burden to undergo pre-planned catheter ablation procedure. **(a)** Engraftment arrhythmia burden of PSC-CM + CA #2 following cell transplantation. **(b)** Bar plot depicting proportion of arrhythmogenic SIRPA^+^CD90^-^CD200^+^ and non-arrhythmogenic SIRPA^+^CD90^-^CD200^-^ cardiomyocytes for each subject in catheter ablation treatment group. **(c)** scRNA-seq analysis of cell dose received by PSC-CM + CA #2.

**Supplementary Figure 11.**
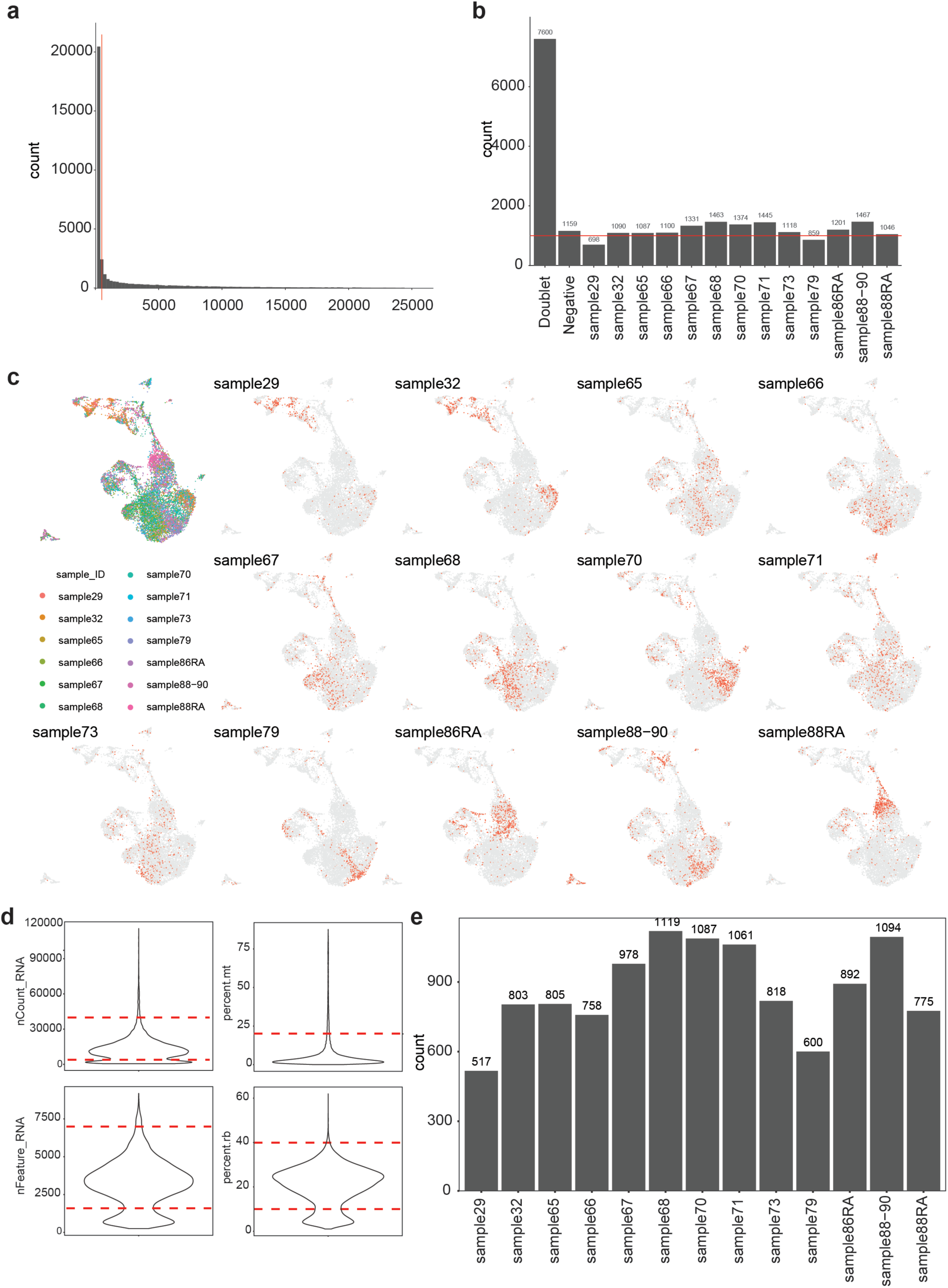
Quality control of scRNA-seq dataset. **(a)** Number of genes against the number of expressing cells. Genes expressed in less than 1% of total cells were removed, and the filter threshold is represented by the red line. **(b)** Samples were demultiplexed and assigned to their original identities or indicated as doublets or negatives. **(c)** UMAP plots of cells after filtering, coloured by the sample of origin. **(d)** Violin plots showing the distribution of quality control metrics including the total number of molecules detected in each cell (nCount_RNA), number of genes detected per cell (nFeature_RNA), percentage of mitochondrial reads (percent.mt), and percentage of ribosome reads (percent.rb). The red dashed line represents the thresholds chosen for filtering. **(e)** Distribution of cells across different samples after quality control filtering.

**Supplementary Table 1.**
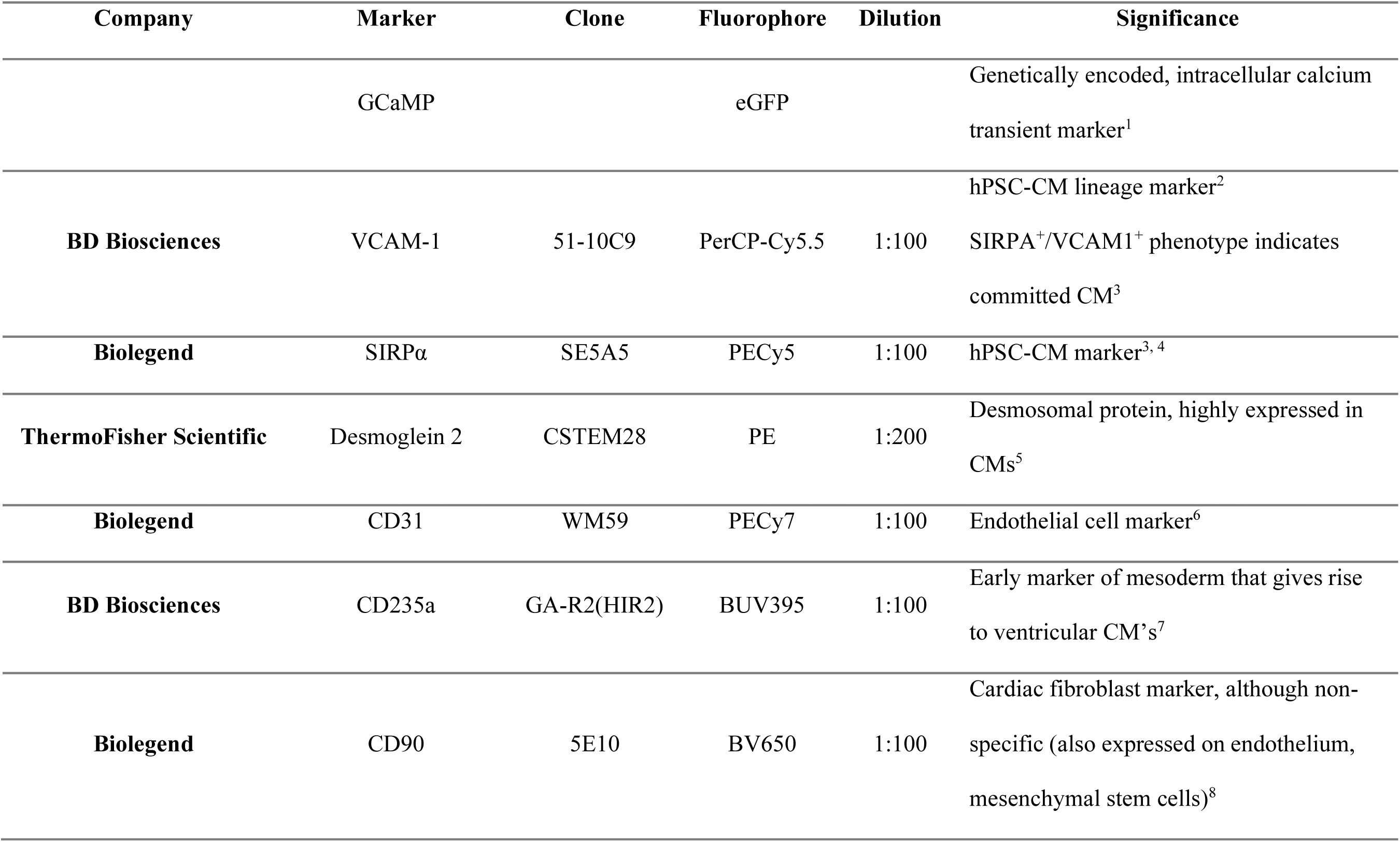
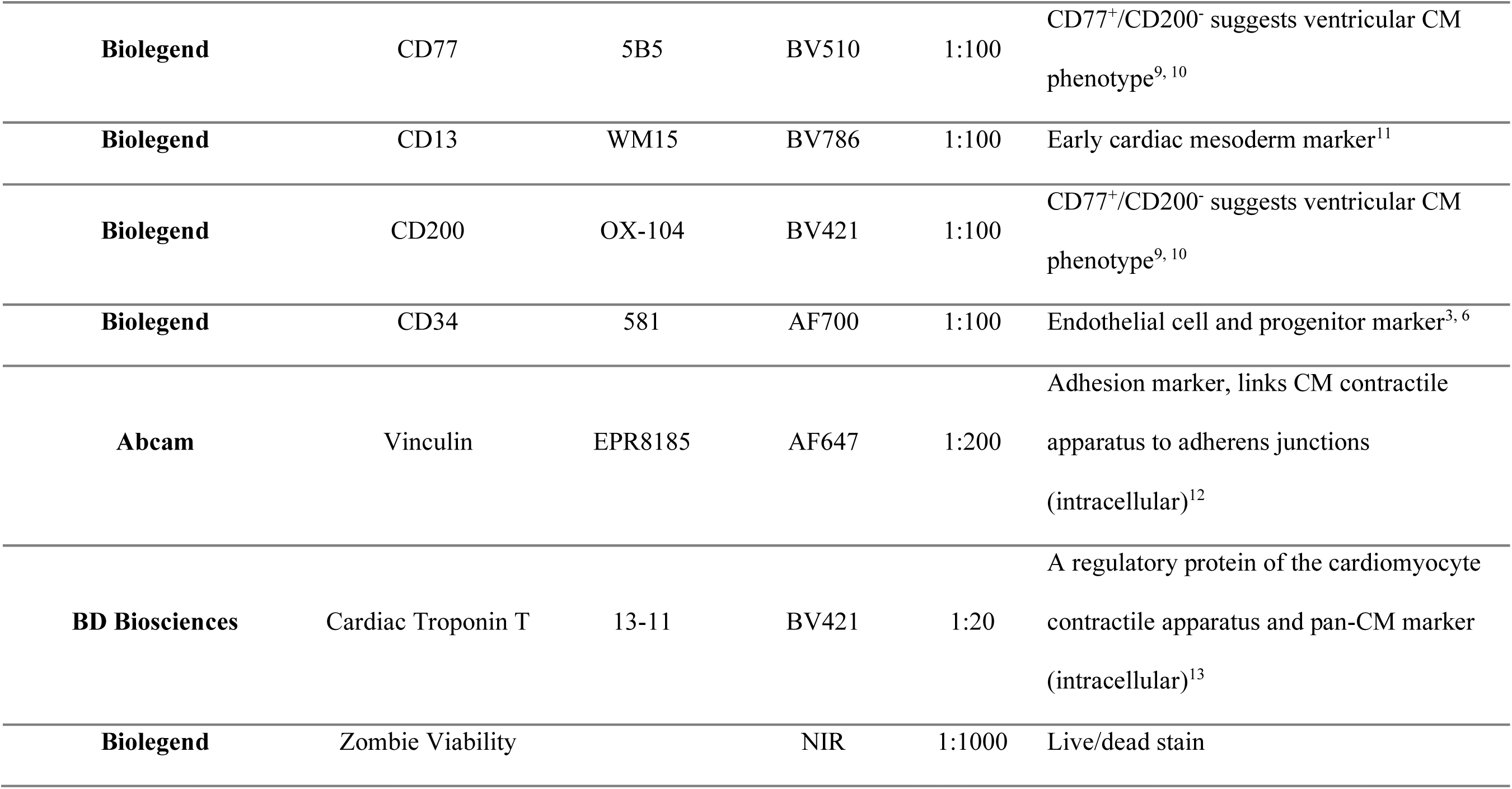
Antibodies used for high-parameter flow cytometry experiments.

**Supplementary Table 2.**
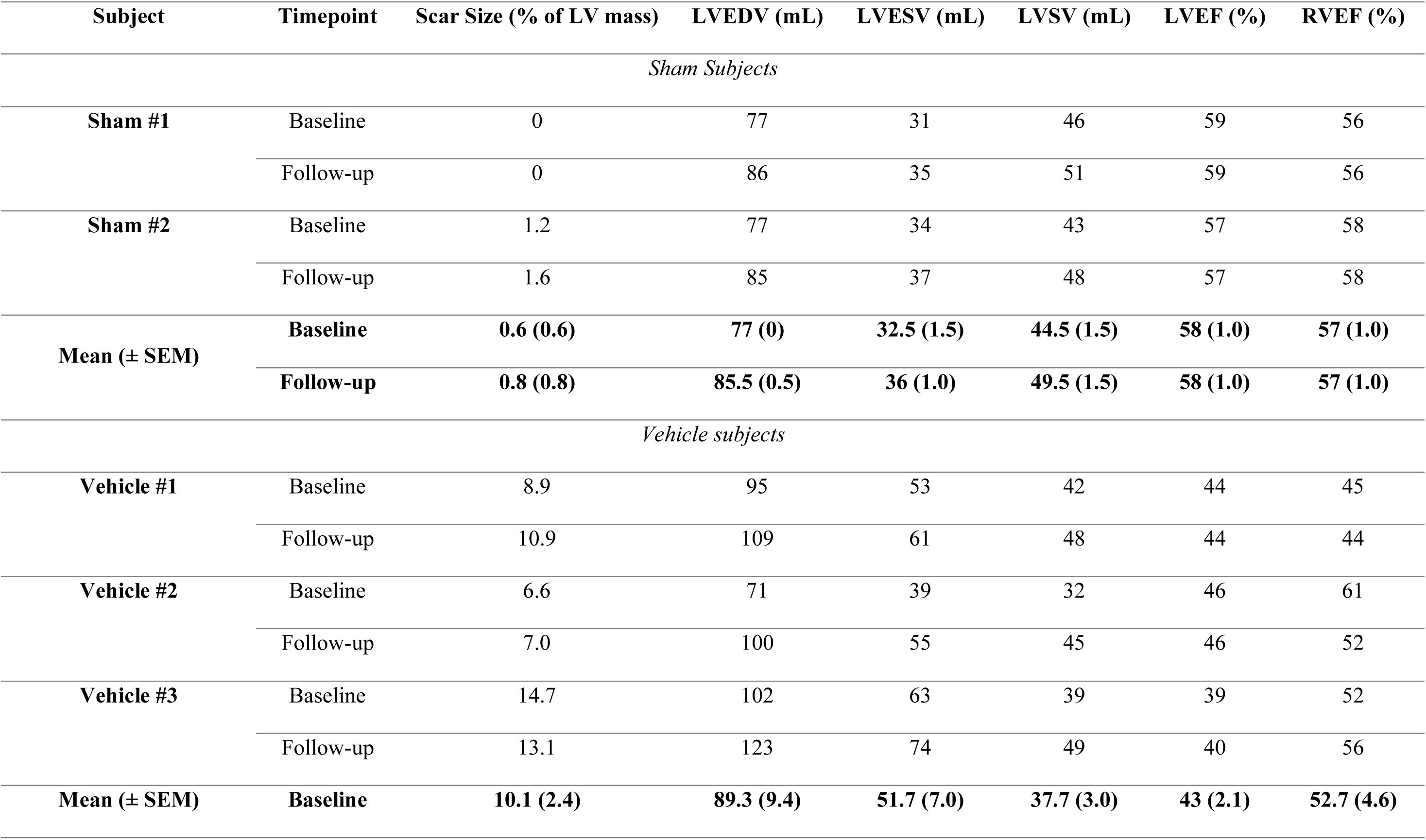
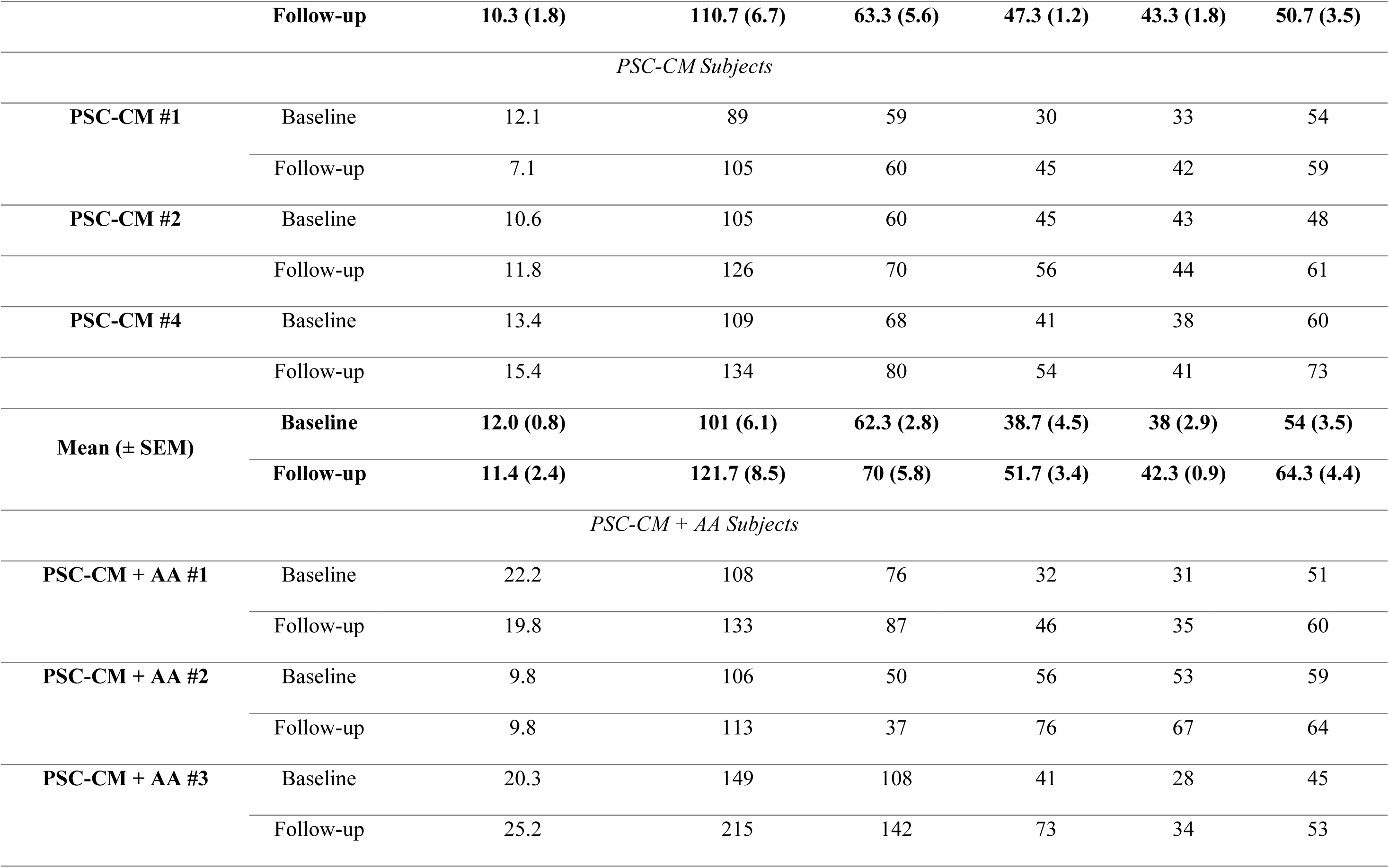
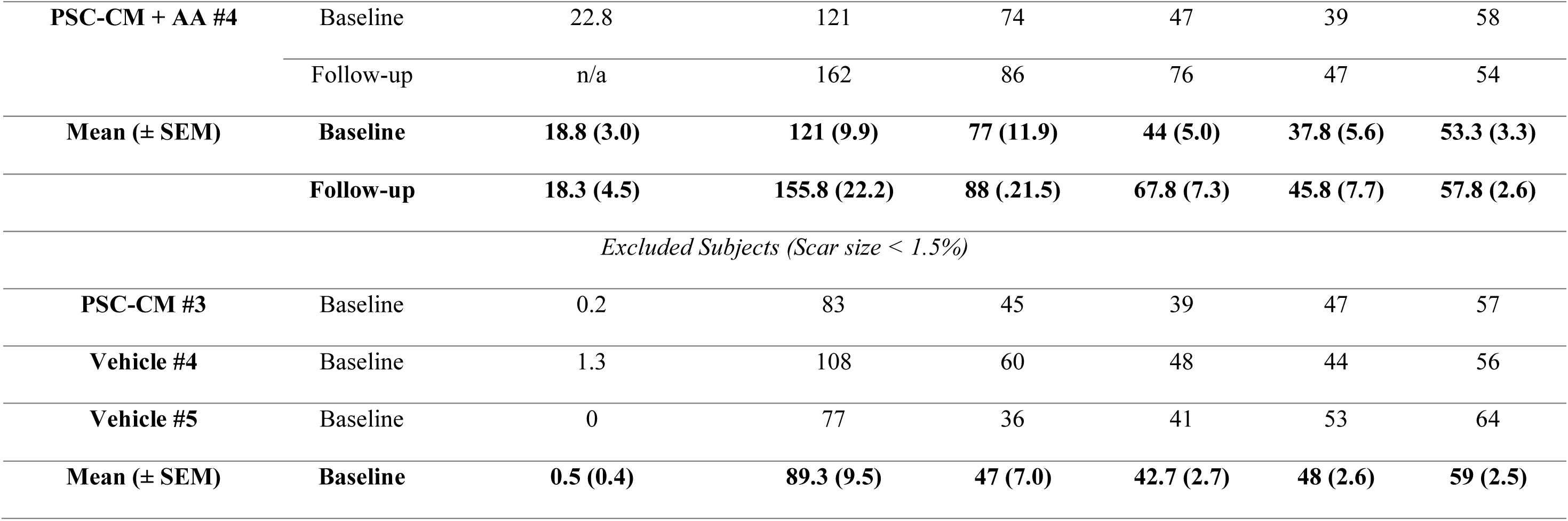
CMR scar size and volumes for Phase 1 large animal experiments.

**Supplementary Table 3.**
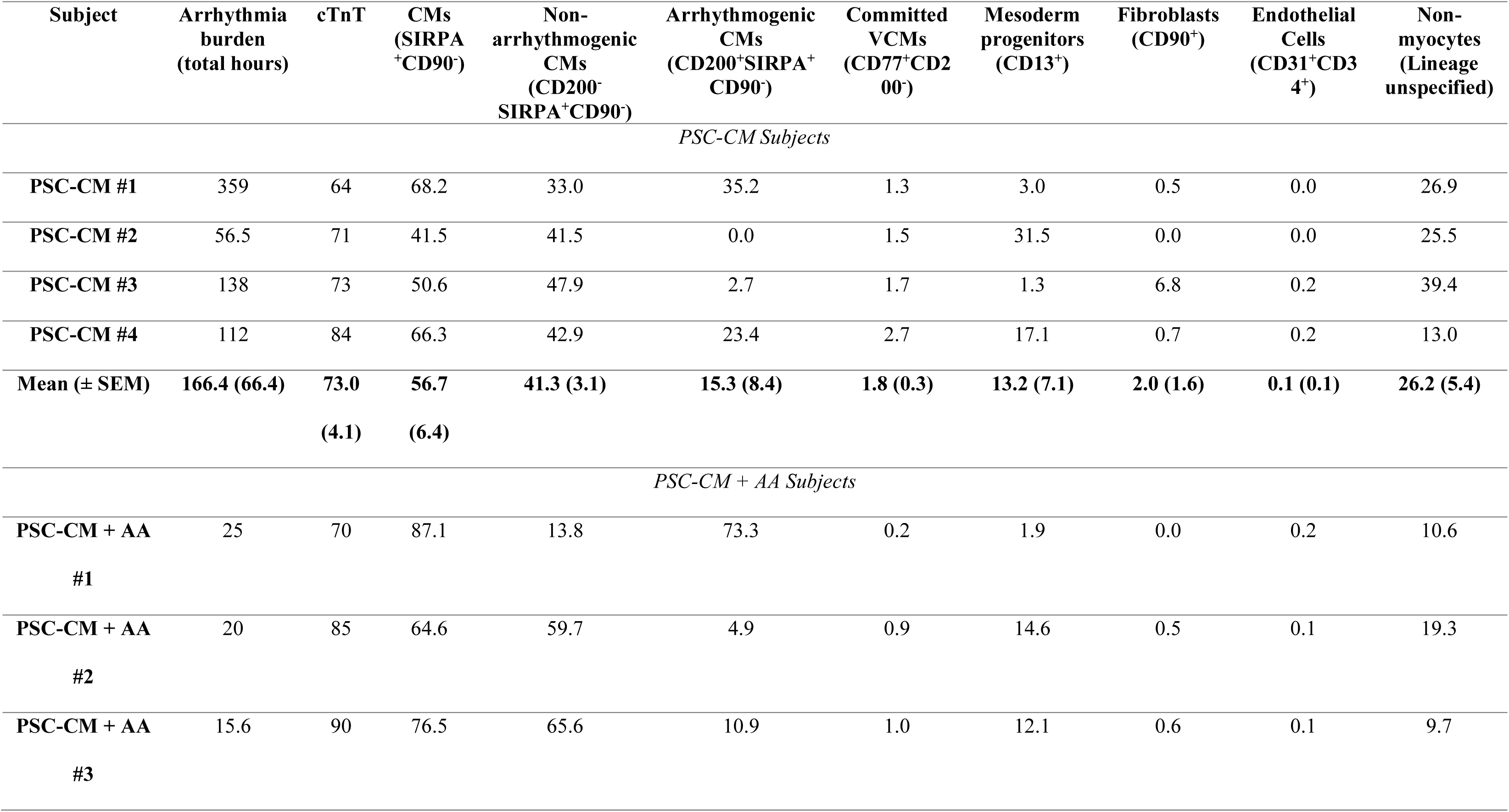
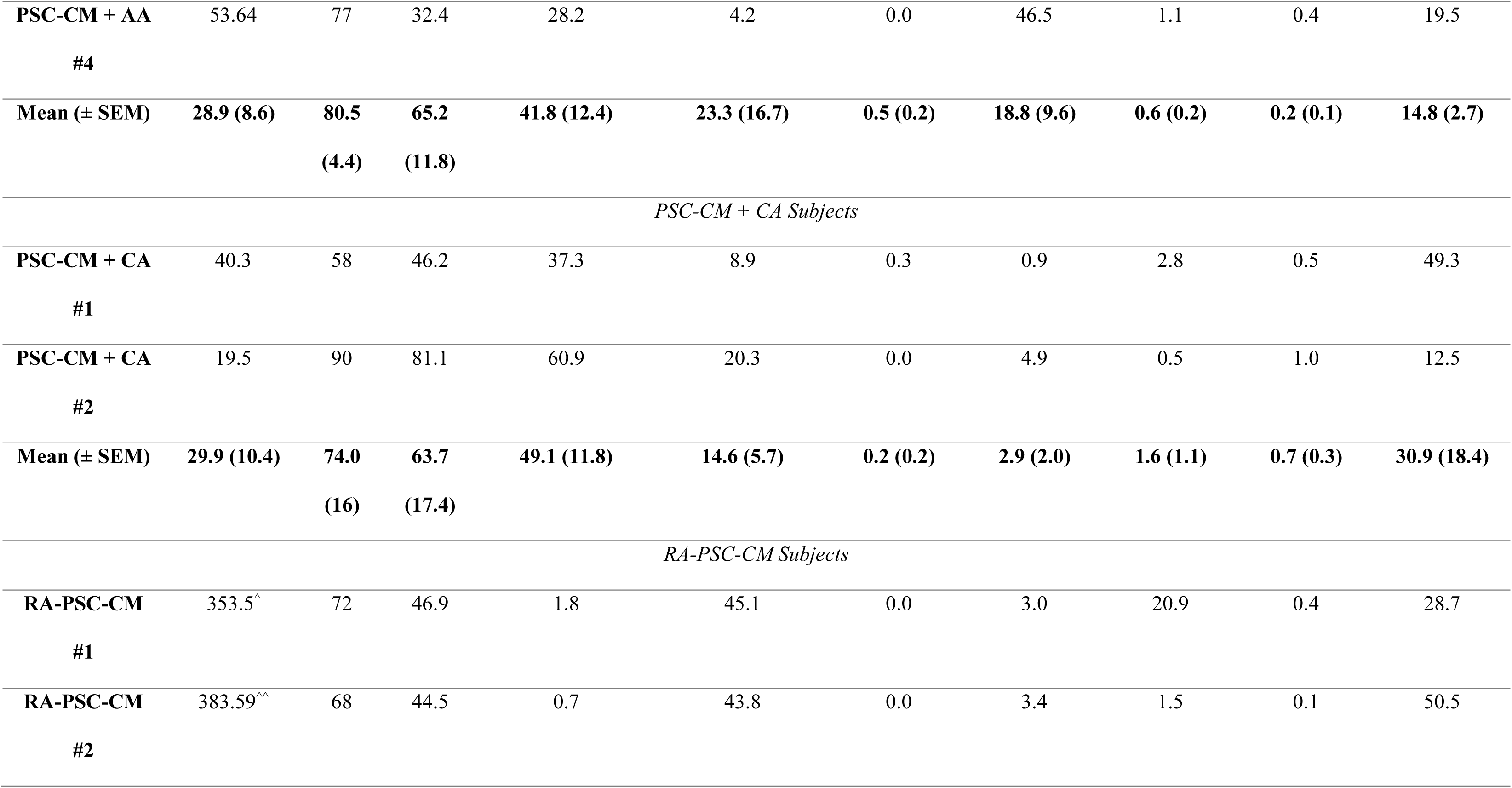
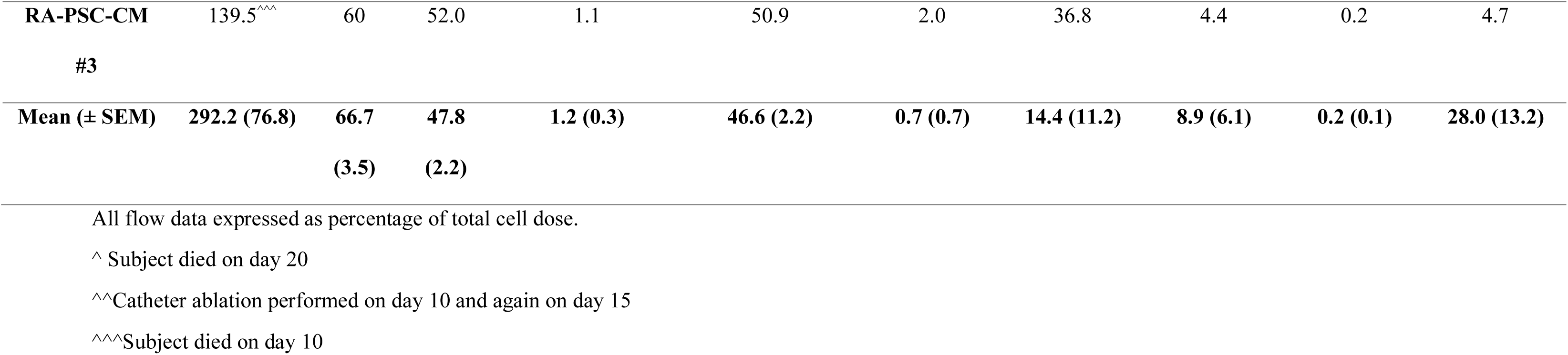
Arrhythmia burden and subpopulation quantification as assessed by high parameter flow cytometry.

**Supplementary Table 4.** Radiofrequency ablation parameters for catheter ablation subjects.

| *PSC-CM + CA #1 (Day 10 post-injection)* |  |  |  |  |  |
| --- | --- | --- | --- | --- | --- |
| **Ablation number** | **Impedance – start ($\Omega$)** | **Impedance – end ($\Omega$)** | **Power (W)** | **Duration (sec)** | **Notes** |
| **1** | 170 | 162 | 30 | 40 |  |
| **2** | 180 | 165 | 30 | 30 |  |
| **3** | 175 | 160 | 30 | 60 | VF induced – 3x 250J shocks required,<br>successfully defibrillated to EA with 3<sup>rd</sup> shock |
| **4** | 165 | 200 | 30 | 24 |  |
| **5** | 170 | 205 | 30 | 18 |  |
| **6** | 170 | 204 | 30 | 24 |  |
| **7** | 170 | 158 | 30 | 30 |  |
| **8** | 168 | 150 | 30 | 21 | Sinus rhythm restored |
| *RA-PSC-CM #3 – Catheter Ablation 1 for EA 1 (Day 10 post-injection)* |  |  |  |  |  |
| **1** | 220 | 237 | 40 | 15 | Impedance cut-off |
| **2** | 190 | 165 | 40 | 30 |  |
| **3** | 200 | 165 | 40 | 30 |  |
| **4** | 220 | 157 | 40 | 30 | VF induced – successfully defibrillated with<br>250J shock back to EA1 |
| **5** | 158 | 242 | 40 | 25 |  |
| **6** | 161 | 259 | 40 | 14 |  |
| **7** | 170 | 246 | 40 | 12 |  |
| **8** | 160 | 249 | 40 | 27 |  |
| **9** | 206 | 197 | 40 | 30 |  |
| **10** | 167 | 240 | 40 | 17 |  |
| **11** | 164 | 247 | 40 | 17 |  |
| **12** | 173 | 156 | 40 | 30 |  |
| **13** | 153 | 158 | 40 | 14 |  |
| **14** | 162 | 156 | 40 | 30 |  |
| **15** | 160 | 149 | 40 | 30 |  |
| **16** | 180 | 154 | 40 | 30 | Sinus rhythm restored |
| *RA-PSC-CM #3 – Catheter Ablation 2 for EA 2 and EA 3 (Day 15 post-injection)* |  |  |  |  |  |
| **1** | 209 | 182 | 40 | 20 |  |
| **2** | 182 | 171 | 40 | 40 | VF induced – successfully defibrillated with<br>250J shock back to EA2 |
| **3** | 237 | 188 | 40 | 40 | Sinus rhythm restored however onset of new<br>EA (EA3) shortly after |
| **4** | 246 | 159 | 40 | 16 | VF induced – successfully defibrillated with<br>250J shock back to EA3 |
| **5** | 167 | 153 | 40 | 40 | Sinus rhythm restored |
| **6** | 160 | 185 | 40 | 19 | Empiric ablation |
| **7** | 160 | 152 | 40 | 38 | Empiric ablation |

**Supplementary Table 5.** Characterisation of working cell bank.

| **Characterisation** | **Method** | **Result** |
| --- | --- | --- |
| **Pluripotency** | Flow cytometry | >90% positive for each of the markers Oct4, Sox2, Tra-1-60 and SSEA4 |
|  | Immunofluorescence | >80% positive for Oct4, Nanog and Tra-1-60 |
| **Mycoplasma** | Lonza Mycoalert Mycoplasma detection assay | Negative |
| **Karyotype** | Stem Cell Technologies Human Pluripotent Stem Cells Genetic Analysis Kit (Product No. #07550) | Negative for 8 common deletions and duplications found in human pluripotent cells |
| **Quantitative evaluation of differentiation capacity to cardiomyocytes** | Suspension based differentiation using small molecules | >80% positive for cardiac troponin at day 15 |
| **Post thaw viability** | N/A | Confluent T75 5 days after thaw of cryopreserved vial. |

**Supplementary Table 6.**
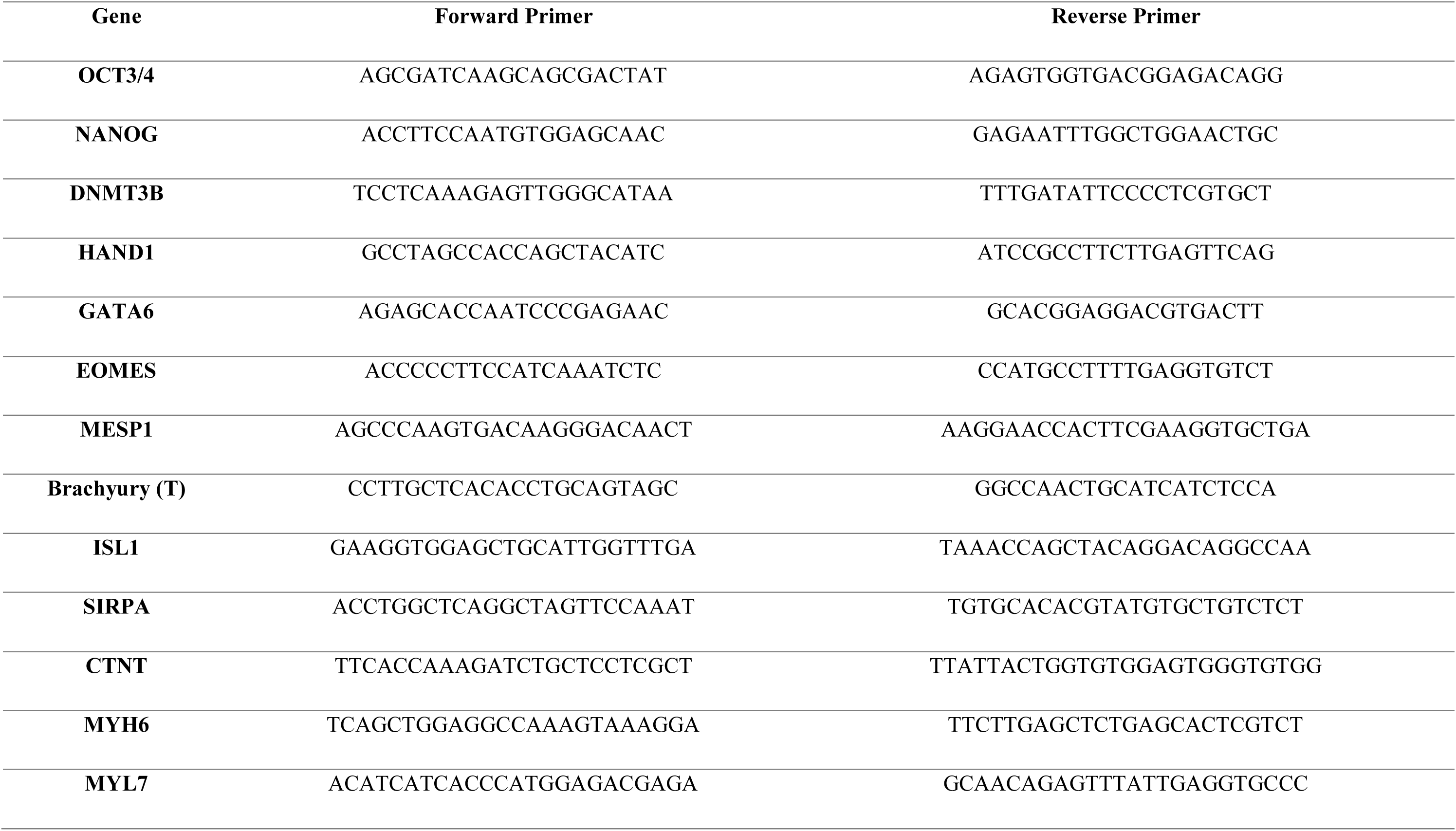
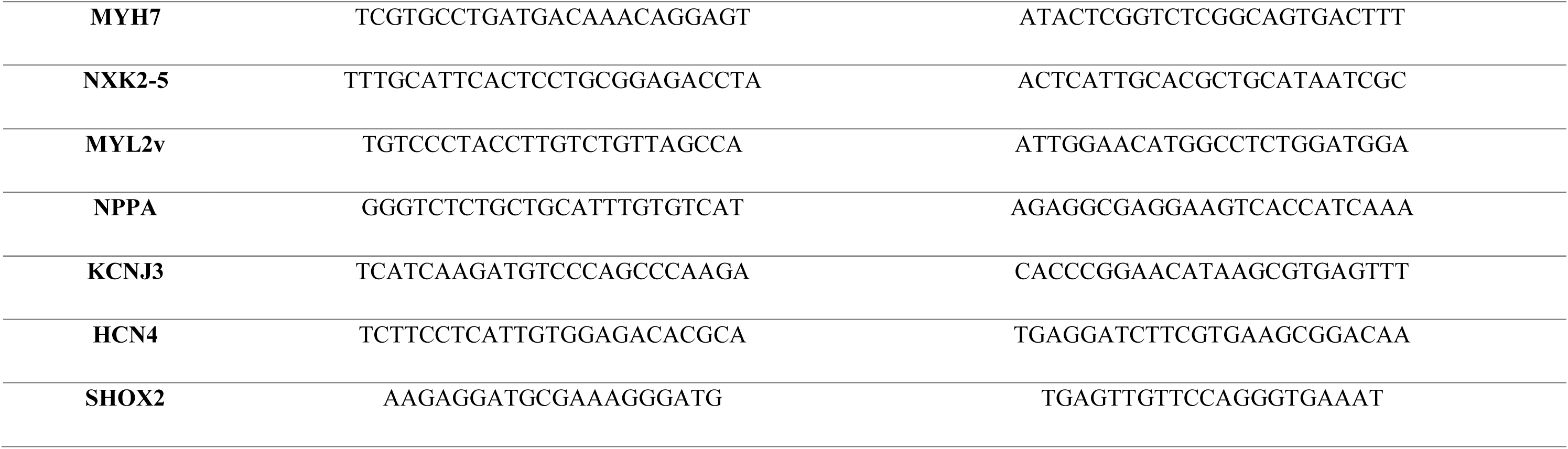
Genes and primers used for qPCR experiments.

**Supplementary Table 7.**
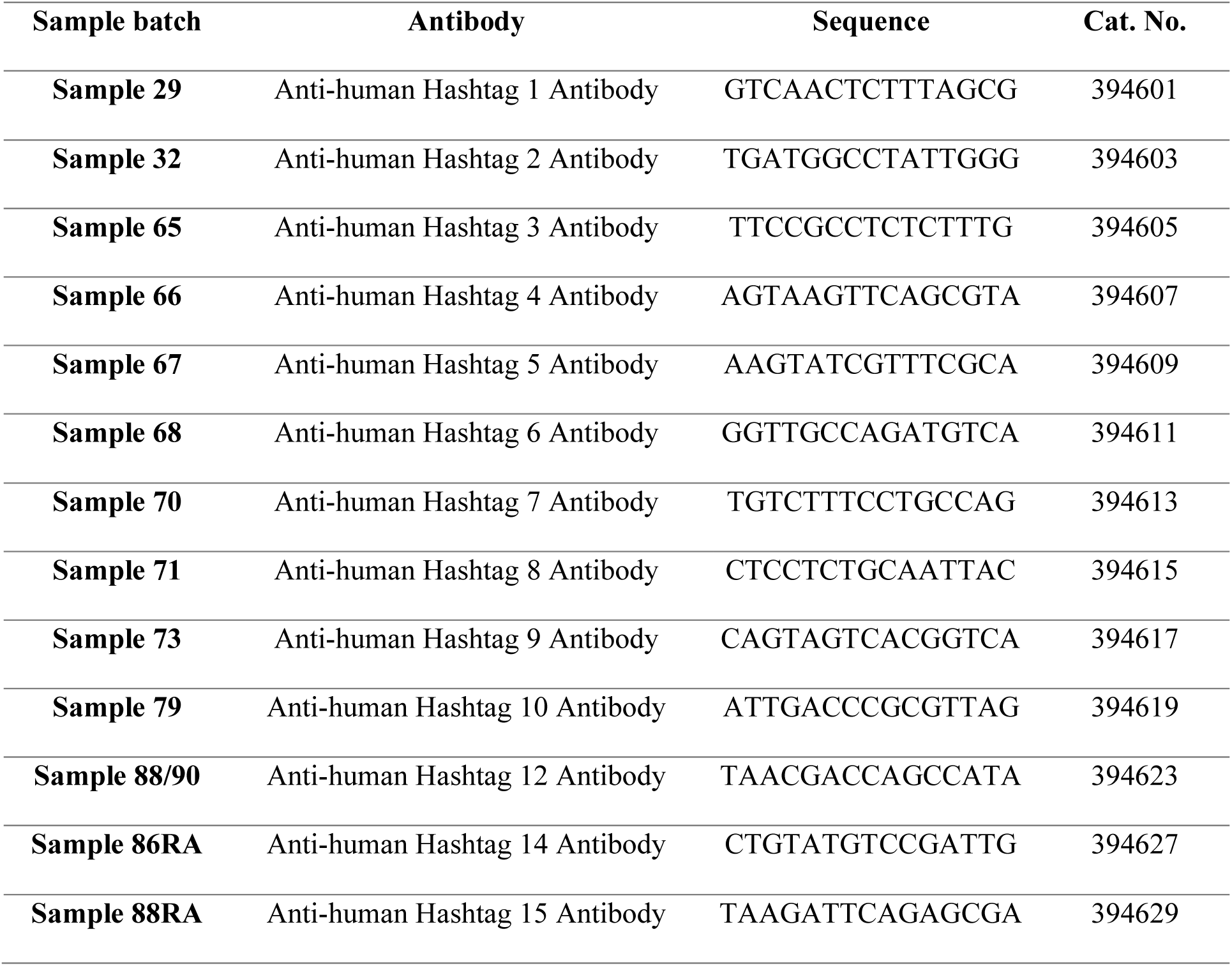
Single cell RNA sequencing hashtag barcode sequences.

**Supplementary Table 8.** Antibodies used for immunohistochemistry experiments.

| **Company** | **Product** | **Marker (Primary)**<br>**Fluorophore**<br>**(Secondary)** | **Dilution** | **Antigen Retrieval**<br>**(Primary)**<br>**Host & Target**<br>**(Secondary)** |
| --- | --- | --- | --- | --- |
| *Primary Antibodies* |  |  |  |  |
| **Dako** | M0851 | Alpha smooth muscle actin | 1:200 | Sodium citrate pH6 |
| **DSHB** | CT3 | Cardiac Troponin T | 2µg/mL | Sodium citrate pH6 |
| **Invitrogen** | PA5-47374 | CD200 | 1:20 | Sodium citrate pH6 |
| **Merck Millipore** | AB1728 | Connexin 43 | 1:100 | Sodium citrate pH6 |
| **Cell Signalling Technology** | C48E7 | Ku80 | 1:500 | Sodium citrate pH6 |
| **Synaptic Systems** | 311011 | MLC-2a | 1:200 | Sodium citrate pH6 |
| **Abcam** | ab79935 | MLC-2v | 1:100 | Sodium citrate pH6 |
|  | ab260039 | SIRPα | 1:100 | Sodium citrate pH6 |
|  | ab28364 | CD31 | 1:50 | Sodium citrate pH6 |
|  | ab13970 | GFP | 1:100 | Sodium citrate pH6 |
| *Secondary Antibodies* |  |  |  |  |
| **ThermoFisher** | A-11029 | Alexa Fluor 488 | 1:500 | Goat anti-mouse |
|  | A-11034 | Alexa Fluor 488 | 1:500 | Goat anti-rabbit |
|  | A-21245 | Alexa Fluor 594 | 1:500 | Goat anti-rabbit |
|  | A-21245 | Alexa Fluor 647 | 1:500 | Goat anti-rabbit |
|  | A-21236 | Alexa Fluor 647 | 1:500 | Goat anti-mouse |
| **Abcam** | ab150173 | Alexa Fluor 488 | 1:500 | Goat anti-chicken |
|  | ab150176 | Alexa Fluor 594 | 1:500 | Goat anti-chicken |
| **Life Technologies** | A11058 | Alexa Fluor 594 | 1:500 | Donkey anti-goat |

## Methods

### Cell Production

#### Bioreactor differentiation protocol

The H9 cell line containing the gCaMP6f fluorescent calcium reporter was used for all experiments (provided by University of Queensland StemCore facility). Use of this cell line and these studies were approved by the University of Queensland Human Research Ethics Committee (Approval numbers: HE002069 & HE000750). Each production run was started from a frozen WCB cryovial, which was expanded for 8 days in monolayer on matrigel (Corning) using the commercially available medium mTeSR^TM^ Plus (Stem Cell Technologies) (Supplementary Fig 1a). The WCB was characterised for pluripotent markers, genetic stability, and differentiation capacity (Supplementary Table 5 & 6). On Day -3, cells were inoculated at a density of 2.5-3.0 × 10^5^ cells/mL in mTeSR 3D supplemented with 10 µM Y-27632 (TOCRIS Bioscience) and 10µg/mL of a thermoresponsive polymer pNIPAM conjugated to a fragment of recombinant fibronectin^31^ to form aggregates. A DASbox (Eppendorf) stirred tank bioreactor system was used for aggregate formation and suspensions controlled at 37.2°C, pH 7.2, and 30% DO. At day -1, rapamycin (Merck) was added to a final concentration of 5nM to enhance survival during differentiation^68^. On day 0, pluripotent aggregates were washed twice with RPMI 1640 then cultured in RPMI-B27 without insulin (Thermo Fisher Scientific) containing 6µM CHIR 99021 (TOCRIS Bioscience) and 5nM of Rapamycin. On day 1, 24 hours post mesoderm induction, aggregates were washed once with RPMI 1640 and returned to RPMI-B27 without insulin, containing 5nM Rapamycin. On day 2, aggregates were washed and returned to RPMI-B27 without insulin, containing 2µM IWP-2 (Stem Cell Technologies). On day 4, aggregates were transferred to RPMI-B27 with insulin and media exchanged every other day until cryopreservation at day 15. Prior to cryopreservation, H9-gCaMP6f derived cardiomyocytes were heat-shocked for 30 minutes at 42°C and treated with a pro-survival cocktail (100ng/mL IGF-1, 0.6µM Cyclosporin A) to enhance their survival after transplantation^13, 14, 69^. Cardiomyocyte aggregates were incubated for 6 hours in 4mg/mL Collagenase IV, washed, and dissociated to single cells using TrypLE (Thermo Fisher). Cardiomyocytes were cryopreserved at 10 × 10^6^ cells/mL in CryoStor CS10 (Stem Cell Technologies).

#### Flow Cytometry

The expression of cTnT was measured on a Cytoflex Flow Cytometer (Beckman Coulter). Briefly, 1 × 10^6^ cells were fixed with 2% paraformaldehyde for 10 min at room temperature. Cells were stored in FACS wash buffer (0.5% BSA in PBS) at 4 °C until staining. Samples were permeabilised in 0.1% Triton X-100 for 10 min then stained with CTNT-FITC 1:50 (Miltenyi Biotec) for 30 min at room temperature. Cells were washed twice before analysis. The excitation laser and emission filters used were: Ex. 488, Em. 525/40.

#### Quantitative PCR

After dissociation of aggregates, 1 × 10^6^ cells were collected in RNAprotect Cell Reagent (Qiagen) and stored at 4 °C until RNA extraction. RNA was extracted using the Qiagen RNeasy Mini kit according to manufacturer’s instructions. Using RevertAid First Strand cDNA Synthesis Kit (ThermoFisher Scientific), 1 µg of RNA was converted to cDNA in a 20 µL reaction according to manufacturer’s instructions. cDNA was diluted 1:10 with DNAse/RNAse free water and 1 µL of diluted cDNA used in each 10 µL reaction with 4 µL of 1µM FIR primers and 5 µL Fast SYBR Green qPCR master mix (ThermoFisher Scientific). Genes and respective primers used are outlined in Supplementary Table 6. Reactions were performed in technical triplicate. Quantitative PCR was performed on the BioRad CFX96 Real-Time PCR Detection System with standard cycling parameters described in the master mix protocol. Dissociation curves were acquired at the conclusion of each run. Fold difference in expression was calculated using the comparative C_T_ formula, using GAPDH cDNA from a reference sample. Depending on the gene and time point, undifferentiated hPSCs or fetal heart RNA were used as the reference sample.

### Single-Cell RNA Sequencing

#### Hashtagging and Sequencing protocol

Frozen PSC-CMs were thawed in the water bath and the content of each vial was transferred to a 50 mL Falcon tube containing RPMI-B27 medium with insulin (ThermoFisher Scientific), 5% FBS (ThermoFisher Scientific), and 10 uM ROCK inhibitor Y-27632 (STEMCELL Technologies). Each sample was spun down at 1000 rpm for 5 mins and the cell pellet was resuspended with filtered PBS plus 2% BSA. Cells were then incubated at room temperature for 10 mins in blocking buffer containing 5uL of Human TruStain FcX solution (BioLegend) plus 100uL of staining buffer (2% BSA + 0.01% Tween20 in filtered PBS). Next, 1uL of hashtag antibody (BioLegend) was added to each sample and incubated on ice for 20 mins. The hashtagging protocol employed allowed for multiple samples to be pooled together in a single sequencing run. The barcode sequences for each hashtag antibody are listed in Supplementary Table 7. Cells were then washed twice with staining buffer and 4 x 10^5^ cells from each stained sample were collected and pooled into a 2 mL Eppendorf tube. Prior to sequencing, a viability test suggested an 80% viability of the captured cells. The cell suspensions were loaded onto 10x Genomics Single Cell 3’ Chips to form single cell gel beads in emulsion (GEMs). The Chromium library was generated and sequenced on a NextSeq 500 instrument (Illumina. The sequencing data were further processed to generate FASTQ files and the raw count matrix using the CellRanger pipeline at the Sequencing Facility at the University of Queensland. The multiplexed samples were demultiplexed to their original identities by mapping of sample reads to GRCh38-3.0.0 human reference genome and hashing sequences.

#### Bioinformatics Analysis

The gene expression count matrix was loaded in Seurat. Data quality control steps are outlined below and summarised in Supplementary Fig. 11. We removed genes that were expressed in less than 1% of total cells prior to creating the Seurat object. The hashtag matrix was added to Seurat object as a new assay separate from RNA. RNA data was normalised with log normalisation and the cell hashing data was normalised with centred log-ratio (CLR) transformation. Following normalisation, cells were demultiplexed and mapped to their original sample identities or assigned as doublets or negatives based on the Seurat HTODemux algorithm together with single cell doublet scoring (scds). We further filtered the cells following the standard quality control (QC) workflow in Seurat. We removed low-quality cells after pre-processing the data and retained 11307 cells for continued analysis. After normalisation and scaling of the data, we performed dimensional reduction and selected 50 principal components (PC) dimensions as input to the unsupervised clustering and uniform manifold approximation and projection (UMAP) plot. Cell type demarcating each cluster was annotated by looking into the expression level of marker genes. Nebulosa package was used for enhanced visualisation of marker genes expression.

### High Parameter Flow Cytometry

#### Antibody staining

Thawed PSC-CMs (∼1.5x10^6^ cells per sample tube) were stained with an amine reactive viability dye (Zombie NIR, Biolegend) for 30 minutes in PBS. The samples were washed twice in PBS + 2% FBS and centrifuged at 225G each time. The samples were then stained with fluorochrome-conjugated membrane marker antibodies (Supplementary Table 1) for 30 minutes in PBS + 2% FBS and washed twice.

For measurement of post-thaw cardiac Troponin T positive cells, separate samples of PSC-CMs were fixed for 20 minutes with 4% PFA and permeabilised in PBS + 0.5% Tween-20. The cells were incubated with anti-human Troponin T antibody in permeabilization buffer for 30 minutes then washed twice in PBS + 2% FBS.

Single colour compensation controls were created using compensation beads as follows: CompBead Plus, BD, for mouse antibodies; AbC Total Antibody, ThermoFisher, for rabbit antibodies; ArC amine reactive beads, ThermoFisher, for Zombie NIR and unstained PSC-CMs for endogenous gCaMP/GFP.

#### Data acquisition and analysis

PSC-CMs and compensation controls were analysed with a BD FACSymphony A5 cytometer and FACS Diva software, using application settings.

Data analysis was performed using FlowJo software (Treestar, Version 10.7.2). An overview of the manual gating and data processing method is shown in Supplementary Fig. 12. Compensated data was manually gated to remove debris, non-viable cells, and doublets. To identify PSC-CM sub-populations based on surface marker expression, dimensionality reduction was performed using t-distributed stochastic neighbour embedding (tSNE), and unsupervised clustering was performed using FlowSOM^70, 71^ (opt-SNE parameters: Gradient algorithm - Barnes-Hut, Learning configuration - opt-SNE, KNN algorithm - exact vantage point tree, Iterations - 1000, Perplexity - 30; FlowSOM parameters: Number of meta-clusters - 25, SOM grid size - 10x10, Node scale - 100%, Set Seed - 3). To compare dose composition between animals and identify potentially pro-arrhythmogenic sub-populations, we down- sampled and concatenated the data prior to dimensionality reduction and clustering.

### *In vitro* Electrophysiology

#### iPSC-CM monolayer differentiation

iPSC-CMs (SCVI-8, Stanford Cardiovascular Institute) were used for in-vitro electrophysiology experiments and were differentiated in 2D monolayer culture. Two days prior to differentiation, PSCs were seeded into Matrigel-coated six-well culture plates, at a density of 1.5x10^6^ cells per well, in mTeSR Plus (STEMCELL Technologies) + 10µM Y-27632. On the first day of differentiation (day 0), media was changed from mTeSR Plus to RPMI 1640 + B27 without insulin + Glutamax + Penicillin/Streptomycin + 6µM of the GS3K inhibitor, CHIR-99021 (Tocris Bioscience). After 24h, the cells were washed with PBS to remove CHIR-99021 and the media was switched to RPMI 1640 + B27 without insulin + Glutamax + Penicillin/Streptomycin. On day 3, the media was replenished and supplemented with Wnt signalling inhibitor, IWP-2 (5µM). On day 5, media was changed to RPMI 1640 + B27 without insulin + Glutamax + Penicillin/Streptomycin. For atrial/pacemaker cell enrichment differentiations, 1µM retinoic acid (Sigma-Aldrich) was added at media change on day 3 and again on day 5. On day 7, media was changed to RPMI 1640 + B27 with insulin + Glutamax + Penicillin/Streptomycin. Following this, media was changed every 2-3 days. Cardiomyocyte beating commenced between days 6 and 7.

#### Patch-clamping

For electrophysiological measurements using high-throughput patch clamping, beating iPSC-CMs were dissociated with TrypLE (ThermoFisher) on day 15 and replated onto Matrigel-coated 12 well plates at a density of 1.5x10^6^ cells per well and maintained in culture until being used for patch-clamping experiments between days 30-35. On the day of patch clamp recording, the cells were washed with PBS, dissociated with TrypLE, and centrifuged for 3 minutes at 300G. The cells were then resuspended in a solution of 80% RPMI-1640 without phenol red + Ca^2+^ + 10%FBS and 20% divalent cation free buffer containing 140 NaCl, 4KCl, 5 Glucose, 10 HEPES, pH7.4 with NaOH. The cells were counted and diluted to a concentration of ∼2x10^5^ cells/mL using the divalent free solution.

Patch clamp recordings were collected using a Syncropatch 384PE (Nanion Technologies), in voltage clamp mode. Sodium current (I_Na_) was recorded with internal and external solutions containing the following (mM): 110 CsF, 10 NaCl, 10 CsCl, 10 HEPES, 10 EGTA ph7.2 with CsOH and 140 NaCl, 4 KCl, 5 glucose 10 HEPES 2 CaCl2, 1 MgCl2 pH7.4 with NaOH, respectively. Sodium current was elicited in 5mV voltage steps ranging from -120 to +40mV from a holding potential of -80mV, with a 200ms pre-pulse to -120mV.

#### Optical electrophysiology

Action potentials were recorded using a kinetic imaging cytometer (KIC, Vala Sciences, San Diego, CA. USA) and voltage-sensitive fluorescent indicator (FluoVolt, ThermoFisher). On day 14 of differentiation, beating cardiomyocytes were dissociated with TrypLE (ThermoFisher) and replated onto Matrigel-coated flat-bottomed 96 well plates (Greiner CELLSTAR, Sigma-Aldrich) at a density of 1.5x10^4^ cells per well. The cells were maintained for a further 5-7 days. On the day of recording, the cells were washed with 100µL RPMI-1640 without phenol red + Ca^2+^. The media was removed and then replaced with 50µL fresh RPMI without phenol red + Ca^2+^ and incubated at 37°C for 60 minutes. The cells were then incubated for 20 minutes in RPMI-1640 without phenol red supplemented with FluoVolt, Powerload pluronic solution, and Hoescht dye. The FluoVolt solution was then removed and replaced with fresh RPMI-1640 without phenol red + Ca^2+^.

The cells were recorded on the KIC for 5 seconds unstimulated, followed by 10 seconds stimulation at 1.5Hz, followed by another 5 seconds unstimulated. Cell segmentation and single cell transient extraction was performed with CyteSeer Scanner software (Vala Sciences, San Diego, CA. USA). The optical action potential measurements were analysed using custom KIC data analysis software (KIC DAT)^72^.

### Porcine Experiments

#### Acclimatation/Housing

All experiments were conducted in female landrace swine (2 to 4 months, 25 - 30kg) acquired from the same local source. All procedures in this study were approved by the Western Sydney Local Health District Animal Ethics Committee (protocol ID: 4262.03.17). Animals were housed in a purpose-built large animal research facility to which they were brought 1-2 weeks prior to the first scheduled procedure for acclimatization.

#### Myocardial Infarction

Myocardial infarction was percutaneously induced by methods previously described^73^. Briefly, animals were premedicated with intramuscular ketamine (10 mg/kg), methadone (0.3 mg/kg), and midazolam (0.3 mg/kg), intubated, ventilated, and maintained with inhalational isoflurane. The left coronary artery was engaged percutaneously via the right femoral artery using a 6F hockey stick guiding catheter (Medtronic, Minnesota, U.S.A). A 0.36-mm coronary guidewire (Sion Blue, Asahi Intecc Co. Ltd., Aichi, Japan) was delivered into the left anterior descending artery (LAD). Myocardial infarction was induced by occluding the mid LAD distal to the first diagonal branch with an inflated 2.0-3.0-mm angioplasty balloon (Boston Scientific, Massachusetts, U.S.A) for 90 minutes. Coronary angiography performed after reperfusion confirmed vessel patency and resolution of ST elevation. Ventricular arrythmias were treated with anti-arrhythmics and defibrillation as required.

#### Telemetry device implantation

All animals underwent telemetry transmitter implantation (easyTEL+, Emka technologies) under general anaesthesia. The transmitter was implanted in a subcutaneous pocket which was created in the left flank. Subcutaneous leads were tunnelled to capture the cardiac apex and base. Signal quality was assessed prior to securing device and leads in situ.

#### Telemetry Analysis

Telemetric ECG and accelerometer data was continuously monitored from the time of device implantation. Semi-automated quantification of heart rate, arrhythmia burden and accelerometer data was performed offline by a cardiologist using the EcgAUTO 3.5.5.16 software package (Emka technologies, Paris, France). Arrhythmia was defined as an ectopic beat (e.g. premature ventricular contraction) or rhythm. The entire recorded dataset for each subject was analysed, with data presented as daily averages (mean ± S.E.M).

#### Cardiac Magnetic Resonance Imaging

Animals were premedicated with intramuscular ketamine (10 mg/kg), methadone (0.3 mg/kg), and midazolam (0.3 mg/kg), before intubation and mechanical ventilation. General anesthesia was induced with intravenous propofol (2-5 mg/kg) and maintained with 2% inhaled isoflurane. Breathing was held in end-expiration for all image acquisitions. All CMR examinations were performed on a Siemens 3T Prisma system (Siemens Medical Systems) utilising an 18-channel body array together with spine array coils and 4-lead electrocardiogram (ECG) gating. Axial and coronal TrueFISP (true fast imaging with steady state precession) sequences through the heart were acquired to plan preliminary 2-chamber, short-axis, and 4-chamber single-slice gradient images. True 2-chamber, 3-chamber, 4-chamber, and right ventricular outflow tract 8mm single-slice TRUFI (true fast imaging) cines were then planned and acquired, as well as a short-axis stack of 14 contiguous 8mm slices, starting just beyond the apex and extending into the atrium, planned from 2-and 4-chamber cine images in end-diastole.

TrueFISP imaging was used for all acquisitions with the following parameters: TE (echo time) 1.3 ms, TR (repetition time) at R-R interval of individual animal, FOV (field-of-view) 370 mm, slice thickness 8mm, in-plane resolution 1.4mm x 1.4mm, flip angle 10 degrees, 25 calculated phases.

Late Gadolinium Enhancement (LGE) images were acquired for tissue characterisation using a segmented inversion recovery fast gradient echo sequence in SAX, LVLA and 4-chamber planes. TI (time of inversion) scout was performed at 9 minutes post contrast to select appropriate inversion time (usually around 300ms) for LGE acquisitions which began at around 11 minutes post-injection. The SAX was run with 30 contiguous slices of 4mm thickness, covering the entire heart with in-plane resolution of 1.5mm x 1.5mm whereas for the LVLA and 4-ch sequences, single 8mm thick slices were acquired with in-plane resolution of 1.4mm x 1.4mm. Other LGE parameters were TR: R-R interval of individual animal, TE: 1.65ms, flip angle: 20 degrees.

Image analysis was performed offline using dedicated software (Medis Suite MR 3.2, Medis Medical Imaging, Schuttersveld 9, The Netherlands) by two independent, blinded cardiologists. Volumetric assessment was performed according to the Society for Cardiovascular Magnetic Resonance guidelines^74^. In brief, short axis end-diastolic and end-systolic images were chosen as the maximal and minimal mid-ventricular cross-sectional areas. End-diastolic epicardial and endocardial borders, along with end-systolic endocardial borders were manually traced for each slice. The vendor specific automated contouring algorithm was not used due to suboptimal performance in non-human subjects. Papillary muscles were included in volume and excluded in mass calculations. The difference between end-diastolic and end-systolic endocardial borders represented the left or right ventricular stroke volume, and ejection fraction was calculated as the stroke volume/end-diastolic volume.

Infarct size was quantified using a separate dedicated software (Segment, Medviso, Lund, Sweden). The short stack late gadolinium enhanced images were used for infarct quantification. Endocardial and epicardial borders were manually contoured from LV base to apex. The Segment automated algorithm for scar quantification was then applied by designating the left anterior wall as the infarct territory. This algorithm delineates ares of late gadolinium enhancement as well as areas of microvascular obstruction. Infarct size was expressed in millilitres(mL). Infarct percent was then calculated by dividing infarct size by end diastolic left ventricular mass, as measured in end diastole on the cine short stack acquisitions. All parameters were analysed by two independent, blinded observers with excellent inter-observer variability. Intra-observer variability was determined by having one observer, blinded to previous measurements, repeat measurements on all subjects on two separate occasions, at least two weeks apart (Supplementary Fig. 4).

#### ADAS-3D CMR Reconstructions

Following CMR image acquisition, A separate offline segmentation software, ADAS-3D (Galgos Medical, Barcelona, Spain), was utilized to process 3D reconstructions of the left ventricle (LV) and fibrosis as identified on late gadolinium enhancement (LGE) for integration in to the electroanatomic mapping system (EAM). Complete cardiac anatomy including the coronary arteries were also exported to assist with registration to EAM reconstructions. Endocardial and epicardial borders were delineated in all slices and an assessment of scar was made based on the pixel signal intensity (PSI) between the two borders. Values for the identification of dense fibrosis and border zone regions using the maximum PSI have been previously correlated with low voltage regions and conducting channels on EAM^75, 76^. Fibrosis area was calculated by averaging the PSI of the endocardial to mid myocardial layers and the average PSI of the mid myocardial to epicardial layers^77^. The calculated values were then projected using the trilinear interpolation method to the LV reconstruction^77^ of the endocardial and epicardial surfaces.

#### Thoracotomy, epicardial mapping and cell injection

On day 0, animals were sedated and returned to the operating theatre for thoracotomy and transepicardial injections. Following cannulation and intubation, an arterial line was inserted into the auricular or distal limb artery to enable continuous blood pressure monitoring. A 100 mcg Fentanyl patch was applied, and intravenous antibiotic (Cephazolin, 1 g) was administered. Intercostal nerve block of the 2^nd^ to 6^th^ rib spaces was achieved with Bupivacaine and Lignocaine.

Thoracotomy was performed by making an incision in the left lateral chest wall at the 4^th^/5^th^ intercostal space. The incision was opened under direct vision using self-retaining rib retractors. The pericardium was opened anterolaterally, and the apex and anterior left ventricle were gently exposed by packing beneath the left ventricle with wet gauze. Haemodynamics were carefully monitored and metaraminol boluses were administered as required to maintain the systolic blood pressure greater than 100mmHg. The epicardial surface of the left ventricle was electroanatomically mapped using an electrophysiological mapping catheter (Navistar Thermocool Smarttouch, Biosense Webster). Cardiac MRI was imported to the EAM system and registration was performed by placing the mapping catheter placed at fiducial landmarks under direct visualization. Fiducial landmarks used included the apex, mitral valve annulus and left anterior descending artery. EAM points were acquired on the EAM system with mapping catheter and identical landmarks were selected on the cardiac MRI. An initial registration was performed with landmarks and then a secondary registration with a best fit of all anatomy was applied using the EAM system software (CartoMerge, Biosense Webster)^78^. Confirmation of registration accuracy was reconfirmed with the EAM catheter. Following registration, the electroanatomic voltage map was assessed to identify scar, border and remote zones. Transepicardial injections of either vehicle (8 x 300 µL injections of RPMI B27) or cells (750x10^6^ cells distributed in 8 x 300 µL injections) were then performed under direct vision into infarct and border zones using a 27-gauge insulin syringe. All pigs were treated with transcutaneous Fentanyl patches (100µg/hour) and intravenous buprenorphine 150-300 µg every 8-12 hours for up to 72 hours post-operatively for analgesia.

#### Central Venous Catheter Insertion and Immunosuppression

All subjects received a three-drug immunosuppressive regimen to prevent xenograft rejection. Commencing 5 days prior to cell injections, animals received oral cyclosporine A (10-15mg/kg twice daily) aiming to maintain trough levels greater than 250ng/mL. On day 0, a 2-lumen 5 French central venous catheter (Teleflex) was implanted in the right jugular vein under ultrasound guidance to facilitate ongoing administration of intravenous immunosuppressants and regular blood draws. Following central line placement and prior to cell injection, 500mg Abatacept (CTLA4-Ig, Bristol-Myers Squibb), 30mg/kg methylprednisone and 3-5mg/kg cyclosporine A was administered intravenously. From day 1 post transplantation until euthanasia, subjects received twice daily oral cyclosporine A (10-15mg/kg) to maintain trough levels greater than 250ng/mL, along with 100mg intravenous methylprednsione daily. Another 250mg dose of intravenous Abatacept was given 2 weeks post transplantation. Prophylactic oral amoxicillin/clavulanic acid was given daily to all subjects to prevent central line and thoracotomy infections, and 30mg oral lansoprazole daily was given for gastrointestinal protection.

#### Anti-arrhythmic treatment

The animals randomised to anti-arrhythmic treatment received a 150 mg IV amiodarone bolus at the time of cell injection. From day 1 post transplantation until euthanasia, they were treated with a regimen of 200 mg oral amiodarone twice daily and 10 mg oral ivabradine twice daily.

#### Electrophysiological Study

Prior to euthanasia, all subjects underwent electrophysiological study under general anaesthesia to determine inducibility, mechanism, and electroanatomic origin of any ventricular arrhythmia. An 8.5 Fr Agilis steerable introducer (Abbott Medical) was inserted into the right common femoral vein under ultrasound guidance through which an Advisor HD Grid (Abbott Medical) multi-electrode electrophysiological catheter was advanced to the right ventricle. A 6 Fr decapolar electrophysiological catheter was positioned into the coronary sinus via the right femoral vein. The Ensite Precision EAM system (Abbott Medical) was used to generate a substrate map delineating scar, border and remote zones using bipolar and unipolar voltage cutoffs of 0.5-1.5mV and 3-8.3mV respectively^79, 80^. The left ventricle was then mapped via a retrograde aortic approach. A substrate map was initially created if the subject was in sinus rhythm. Following substrate mapping, VT induction via programmed electrical stimulation (PES) was performed. A drive train of eight beats at 400ms was followed by up to four extra-stimuli delivered one at a time. Initial extrastimuli were delivered at a coupling interval of 300ms which was then decreased by 10ms until ventricular refractoriness. A modified “Michigan” protocol was additionally performed^81^. The Michigan protocol uses exclusively four extra stimuli. At each drive train cycle length of 350ms, programmed stimulation is initiated with coupling intervals of 290, 280, 270, and 260ms for the first through fourth extrastimuli. The coupling intervals of the extra stimuli are shortened simultaneously in 10-ms steps until an extrastimuli is refractory or an arrhythmia is induced. Both PES protocols were repeated at two sites. Following PES an isoprenaline bolus (20mcg) and infusion (6-10mcg/min) was administered with burst pacing starting from 300ms and reduced in 20ms steps until refractory during isoprenaline delivery and washout^82^. The mechanism of any spontaneous or induced ventricular arrhythmias was elucidated through standard pacing maneuvers followed by EAM to identify arrhythmia origin.

#### Catheter Ablation

Subjects who underwent catheter ablation for treatment of EA were premedicated with intramuscular ketamine (10 mg/kg), methadone (0.3 mg/kg), and midazolam (0.3 mg/kg), intubated, ventilated, and maintained with inhalational isoflurane. An arterial line was inserted into the auricular or distal limb artery to enable continuous blood pressure monitoring. All subjects were in spontaneous engraftment arrhythmia at the time of the ablation procedures. Arrhythmia origin was electroanatomically mapped as described above. At site of earliest activation mapping, radiofrequency ablation lesions were delivered using a 4 mm tip open-irrigation catheter (Flexability, Abbott Medical). A single grounding patch was placed on each animal and ablation was performed in power control mode using 30-40W of power and 13 mL/min irrigation flow rate with normal saline. Delivery of each lesion was attempted for 30-60 seconds unless terminated prematurely due to catheter movement or impedance rise. Ablations were terminated after sinus rhythm was restored, followed by an attempt at VT induction as described above.

#### Euthanasia and tissue harvest

Following the terminal EPS procedure, subjects were euthanised with potassium chloride (75- 150mg/kg) under deep anaesthesia and hearts were excised and fixed in 10% neutral buffered formalin for subsequent analysis.

### Histology

#### Tissue Processing

Pig hearts were fixed whole in 10% neutral buffered formalin for a minimum of 48 hours. The ventricles were then sliced into transverse sections of approximately 1cm thickness from the apex (Level 1) towards the base (Level 7). After fixation, the tissues were switched to 70% ethanol, processed and paraffin embedded. Processing, embedding, and sectioning of Level 1 blocks was performed at the Westmead Institute for Medical Research. Processing, embedding, and sectioning of large blocks (Levels 2 and above) was performed by Veterinary Pathology Diagnostics Services at the University of Sydney.

#### Immunohistochemistry

Immunohistochemistry analyses were performed on 4µm sections taken from Levels 1, 2, or 3. Sections were deparaffinised in xylene and hydrated by sequential incubation in 100%, 95%, 70%, 50% ethanol, and water. Antigen retrieval was performed using heated sodium citrate buffer and the sections were then washed in PBS + 0.1% Tween 20, blocked with 5% goat serum in PBS + 0.05% Tween-20, and stained with primary antibodies overnight at 4°C (Supplementary Table 8). The following day, the sections were washed and incubated with secondary antibodies (Supplementary Table 8) for one hour at room temperature, in the dark. The sections were then washed again and incubated with DAPI (1µg/mL, Sigma-Aldrich/Merck) for 10 minutes, rinsed with PBS and mounted with PBS: Glycerol. For demonstration of PSC-CM grafts, and radiofrequency ablation site, whole mount sections from Level 2 tissue blocks were stained with primary antibodies targeting human Ku80, and cardiac troponin T, as described above. For brightfield detection of secondary staining, we used the ImmPRESS Duet double staining polymer kit HRP/AP (Vector Laboratories, MP-7714), and then counterstained sections with aniline blue (1% aniline blue in MilliQ water for 1 minute).

#### Imaging and analysis

Immunofluorescence and brightfield microscopy were performed on an Olympus VS120 Slide Scanner, with the 20X objective (UPLSAPO 20X/ NA 0.75, WD 0.6 / CG Thickness 0.17 or the 40X objective (UPLSAPO 40X/ NA 0.95, WD 0.18 / CG Thickness 0.11-0.23). Images were acquired using Olympus VS-ASW 2.92 software and processed using Olympus VS-DESKTOP 2.9.

Confocal images of GFP, Cx43 and cTnT immunostained pig tissues were taken on a Leica SP8 microscope using tuneable white light and 405nm lasers. Filters were adjusted for optimal signal detection of AlexaFluor488, AlexaFluor594 and AlexaFluor647 using a 2-sequence scanning approach. Z-stacks (7µm with a 0.5mm step size) were imaged using a confocal pinhole set at 1 airy unit

### Statistical analyses

Continuous data are presented as mean ± standard error of mean (SEM). Normality was assessed using the Shapiro-Wilk test, with appropriate parametric or non-parametric tests performed depending on distribution of data. Statistical comparisons of normally distributed data were conducted using unpaired t tests or ordinary one-way ANOVA followed by the Sidak’s post hoc test to adjust for multiple comparisons. Non-normal data was statistically compared using the Mann-Whitney or Kruskal-Wallis test followed by Dunn’s test to adjust for multiple comparisons. Correlations were expressed using Pearson’s or Spearman’s correlation coefficients. Inter- and intra-observer CMR analysis variability was expressed using Bland-Altman plots. Survival analyses were performed using the Kaplan-Meier method, and the log-rank test was applied to determine significance between overall survival between groups. *P* values <0.05 were considered statistically significant. All analyses were performed using GraphPad Prism Version 9.3.1 software.

